# Changes in the genetic requirements for microbial interactions with increasing community complexity

**DOI:** 10.1101/290353

**Authors:** Manon Morin, Emily C. Pierce, Rachel Dutton

## Abstract

Microbial community structure and function rely on complex interactions whose underlying molecular mechanisms are poorly understood. To investigate these interactions in a simple microbiome, we introduced *E. coli* into an experimental community based on a cheese rind and identified the differences in *E. coli’*s genetic requirements for growth in interactive and non-interactive contexts using Random Barcode Transposon Sequencing (RB-TnSeq) and RNASeq. *E. coli’s* genetic requirements varied among pairwise growth conditions and between pairwise and community conditions. Our analysis points to mechanisms by which growth conditions change as a result of increasing community complexity and suggests that growth within a community relies on a combination of pairwise and higher order interactions. Our work provides a framework for using the model organism *E. coli* as a probe to investigate microbial interactions regardless of the genetic tractability of members of the studied ecosystem.

## INTRODUCTION

Microorganisms rarely grow as single isolated species but rather as part of diverse microbial communities. In these communities, bacteria, archaea, protists, viruses and fungi can coexist and perform complex functions impacting biogeochemical cycles and human health (Falkowski, Fenchel, and Delong 2008; Flint et al. 2012). Deciphering microbial growth principles within a community is challenging due to the intricate interactions between microorganisms, and between microorganisms and their environment. While interest in microbial communities has dramatically increased, our understanding of microbial interactions within communities is lagging significantly behind our ability to describe the composition of a given community. Further, the way in which these interactions are organized within a community, such as whether they consist of predominantly pairwise or higher-order interactions, is even less clear. A more precise understanding of microbial interactions, their underlying mechanisms, and how interactions are structured within a community, are all necessary to elucidate the principles by which a community is shaped. In this study, we combine genome-scale genetic and transcriptomic approaches within an experimentally tractable model microbial community to begin to address these questions.

Genome-scale approaches, such as transposon-mutagenesis coupled to next-generation sequencing (TnSeq approaches) have been successfully used to quantify the contribution and thus the importance of individual genes to a given phenotype (van Opijnen and Camilli 2013). Recently, generation and introduction of unique random barcodes into transposon mutant libraries made this approach more high-throughput and less laborious, enabling screens of important genes within hundreds of conditions and for numerous genetically tractable microorganisms (Wetmore et al. 2015; Price et al. 2016). In order to investigate the genetic bases of microbial interactions, we have adapted this approach to allow us to identify and compare genetic requirements in non-interactive and interactive conditions. To do so, we introduced a large and diverse transposon library generated previously in the genetically-tractable model bacterium *E. coli* to characterize the genetic requirements of interactions within a model community based on the rind of cheese (such as Camembert) (Wetmore et al. 2015). We (i) identify the set of important genes for *E. coli* when growing alone in the cheese environment (ii) identify the set of important genes for *E. coli* growth in pairwise conditions with each individual community member and (iii) identify the set of important genes for *E. coli* growth with the complete community. Characterization of the functions or pathways associated with growth in interactive versus non-interactive conditions can then be used to infer the biological process involved in interactions. Additionally, we compared the set of important genes in pairwise conditions with the ones important for growth in a community to investigate how microbial interactions change depending on the complexity of the interactive context.

We further performed two complementary approaches to defining interactions within this system. First, we measured changes in the transcriptional profile of *E. coli* during growth alone, growth in pairwise conditions, and within the community, using RNAseq. This analysis revealed a deep reorganization of gene expression whenever *E. coli* is in the presence of other species. Next, as *E. coli* is a non-endogenous species in our model ecosystem, we generated a transposon library in the endogenous species *Pseudomonas psychrophila* JB418, and performed similar RB-TnSeq analysis during non-interactive and interactive conditions.

This work revealed numerous interactions between species, such as metabolic competition for iron and nitrogen, as well as cross-feeding from fungal partners for certain amino acids. It also highlighted the need for resistance to toxic compounds and osmotic stress as a requirement for growth. Our analysis showed that most of the metabolic interactions (competition and cross-feeding) observed in pairwise conditions are maintained and amplified by addition of partners and similar pairwise interactions. However, around half of the genetic requirements observed in pairwise conditions were no longer apparent in the community, suggesting that higher order interactions emerge in a community.

## RESULTS

### Identification of the basic genetic requirements for growth of the *E. coli* sensor species in isolation

We used the *E. coli* Keio_ML9 RB-TnSeq library from Wetmore et al., 2015, containing 152,018 different insertion mutants (covering 3728 of 4146 protein-coding genes), each associated with a unique 20 nucleotide barcode. This library was originally generated in and maintained on lysogeny broth medium (LB), and was used previously to identify genes required for growth across a variety of conditions (Wetmore et al. 2015; Price et al. 2016). To determine genes important for growth on our cheese-based medium, we grew the library by itself on sterile cheese curd agar plates (CCA: 10% freeze-dried fresh cheese, 3% NaCl, 0.5% xanthan gum, 1.7% agar) (Figure 1A), the same medium used in all further experiments, and used previously to demonstrate that cheese communities could be successfully reconstructed *in vitro* (Wolfe et al. 2014). As genetic requirements are likely to change over the course of growth, we grew the library on CCA for 1, 2 and 3 days. For each time point, we harvested the library from the surface of the cheese plate, extracted genomic DNA, used PCR to amplify the barcoded regions of the transposons, and then sequenced these products to measure the abundance (i.e the number of sequencing reads associated with each barcode) of each transposon mutant over time (see Materials and Methods). For each gene, there are on average 15 individual insertion mutants present in the library. The fitness of each insertion mutant was calculated as the log2 ratio of its abundance at a given timepoint compared to T0. Then, we calculated the weighted average of the fitness of all insertions in a given gene to determine the overall fitness associated with each gene and calculate a corresponding t-score that accounts for the fitness consistency within individual insertions of that gene and thus for the confidence we can attribute to that gene fitness. All gene fitness values were normalized so that a gene with no effect on the growth phenotype has a fitness value of 0 (see (Wetmore et al. 2015) for details).

**Figure 1:**
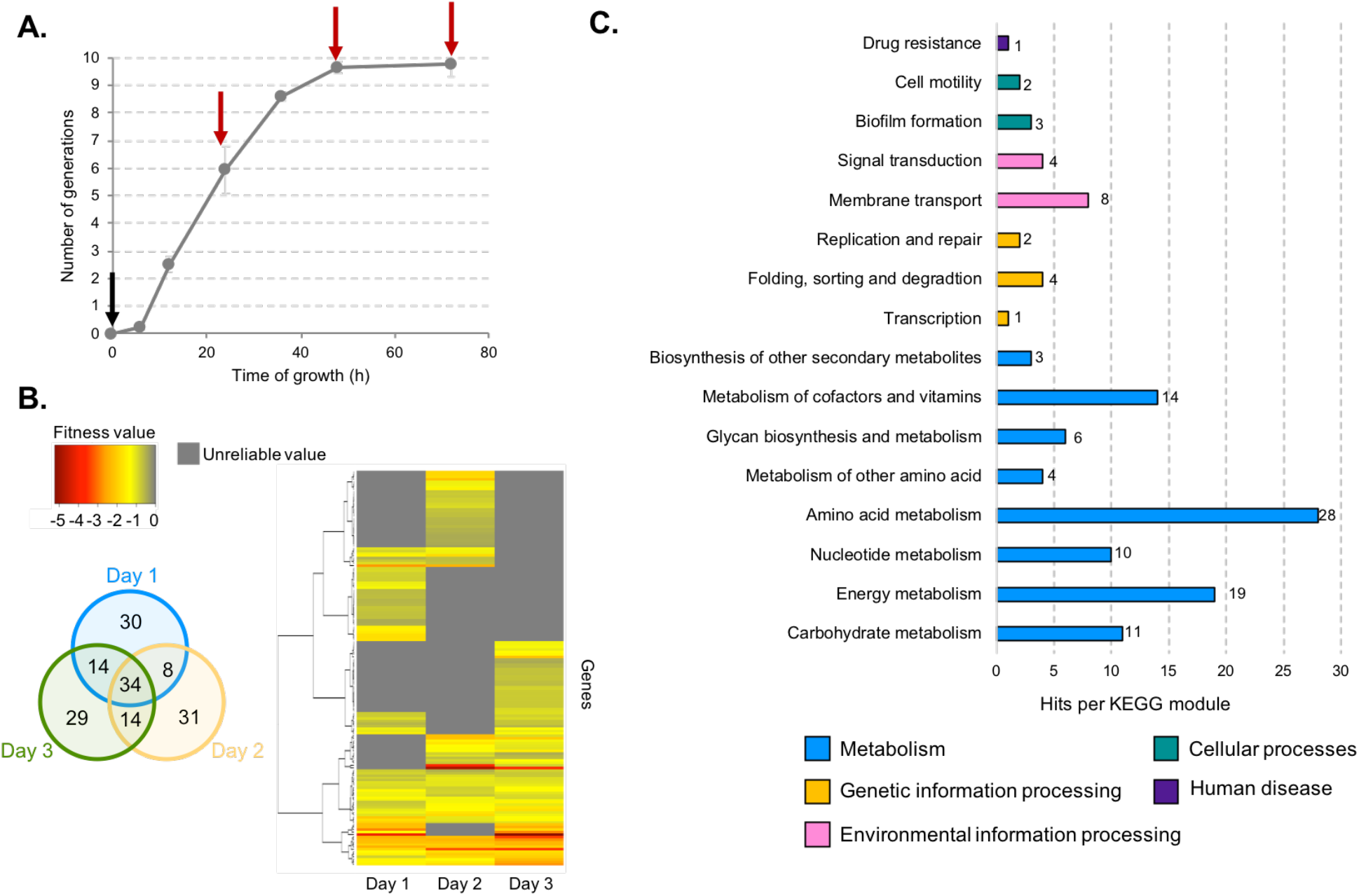
Identification of genes essential for *E. coli* growth alone on cheese curd agar. The E. coli RB-TnSeq library Keio_ML9 (Wetmore et al., 2015) was grown for 3 days on cheese curd agar (CCA). Samples were harvested at T=0h (black arrow), 24h, 48h and 72h (red arrows) for gene fitness determination **(A)**. Genes with a significant negative fitness effect (“essential genes”, abs(t-score) >= 3 and fitness < 0) were identified at day 1 (24h), day 2 (48h) and day 3 (72h) **(B)**. Pooled together, they represented 160 essential genes for E. coli’s growth on CCA. 84 of the 160 genes had hits when mapped to the KEGG BRITE database **(C)**.

At each timepoint, the fitness was calculated for 3298 protein-coding genes. First, we removed genes in which the fitness value did not pass our confidence threshold (absolute t-score >= 3). We then removed genes with positive fitness (day 1 n= 10, day 2 n=14 and day 3 n=16). Thus, we only retained the genes whose deletion leads to a consistent growth defect for *E. coli* on CCA (hereafter referred as to essential genes). This filtering process left 160 genes that were important for *E. coli* growth alone on CCA with a false discovery rate of 0.2% (n=86 for day 1, n=87 for day 2, n= 91 for day 3) (Figure 1B and Supplementary file 1).

To identify the functions that are associated with these 160 genes, we mapped them to the KEGG Brite database (Figure 1C). 84 genes were assigned to KEGG modules and 64 of them were associated with *E. coli* metabolism. Within these metabolic genes, we found 28 genes associated with amino acid metabolism, specifically the biosynthesis of all amino acids except for proline, lysine and histidine. Quantification of free amino acids in our medium highlighted very low concentrations of all amino acids (supplementary figure 1) suggesting that a limited supply of free amino acids leads to a genetic requirement for amino acid biosynthesis. This is supported by the observation that both *spoT* and *relA*, regulators of the stringent response which can be triggered by amino acid starvation (Cashel and Rudd 1996), are also essential. 19 essential genes were associated with energy metabolism and more specifically with sulfur assimilation (n=7 genes) and respiration (n=8 genes). Here, we deduce that essentiality of sulfur assimilation is directly correlated with the lack of the amino acids cysteine and methionine which are the major pools of sulfur-containing compounds in the cell. As a non-endogenous species, *E. coli* might not possess the adequate peptidases or proteases to degrade and use the highly available protein casein. Identification of two of the three genes of the Leloir pathway (*galE* and *galT*), involved in the uptake and conversion of galactose into glucose, suggests that galactose might be a crucial nutrient for *E. coli* growth on CCA. Finally, 8 genes mapped to membrane transport and were associated with two specific pathways: ferric-enterobactin transport and glycine-betaine transport. Ferric-enterobactin transport allows the cells to scavenge iron in a low-iron environment (Raymond, Dertz, and Kim 2003; Hider and Kong 2010). Iron is an essential micronutrient and cheese is known to be iron-limited (Albar et al. 2014). Glycine betaine is used by the cells as an osmoprotectant against high osmolarity environments. During cheese curd processing, high concentrations of NaCl are added (Guinee 2004), and our CCA medium contains 3% NaCl to mimic these conditions. The importance for *E. coli* to maintain its cell osmolarity is also suggested by the essentiality of genes responsible for the transport of the ions sodium, potassium or zinc.

In summary, conditionally essential functions for *E. coli* to grow alone in our experimental environment involved (i) response to low iron availability, (ii) response to osmotic stress and (iii) response to low available nutrients (specifically free amino acids). These required functions are consistent with recently published results about the requirements of the mammary pathogenic *E. coli* (MPEC) during growth in milk (Olson et al. 2017) except for resistance to osmotic stress which does not occur in milk.

To validate the results obtained with the transposon library, we measured the fitness of individual knockout mutants from the *E. coli* Keio collection (Baba et al. 2006). We tested 25 knockout mutants corresponding to genes with a strong growth defect observed after one day of growth. We carried out competitive assays between each knock-out mutant and the wild-type strain on CCA. We calculated each knock-out mutant fitness as the log fold change of its abundance after one day of growth. A z-score was also calculated to assess the confidence of that fitness. 21 of 25 knock-out mutants displayed a fitness value lower than 0 with at least 95% confidence (supplementary figure 2). The remaining 6 mutant strains (*brnQ*, *cysK*, *cysQ*, *serA*, *trxA* and *waaP*) were associated with high fitness value variability across replicate experiments and thus they were associated with a lower z-score. Altogether, this supports the reliability and validity of RB-TnSeq results.

### Identification of genes essential for *E. coli* growth in pairwise conditions

The growth of our *E. coli* library alone allowed us to determine the baseline set of genes required for optimal growth in the model cheese environment. We next wanted to examine the genes required for growth when another species is present. First, we analyzed the growth of *E. coli* and the partner species. We grew *E. coli* for 3 days on CCA in the presence of either *H. alvei*, *G. candidum* or *P. camemberti*. In addition to belonging to distinct domains or phyla, these 3 partners are the typical members of a bloomy rind cheese community (such as Brie or Camembert). The presence of *E. coli* did not influence the growth of any species (supplementary figure 3). However, *E. coli’*s growth was reduced in the presence of each partner (Figure 2A).

**Figure 2:**
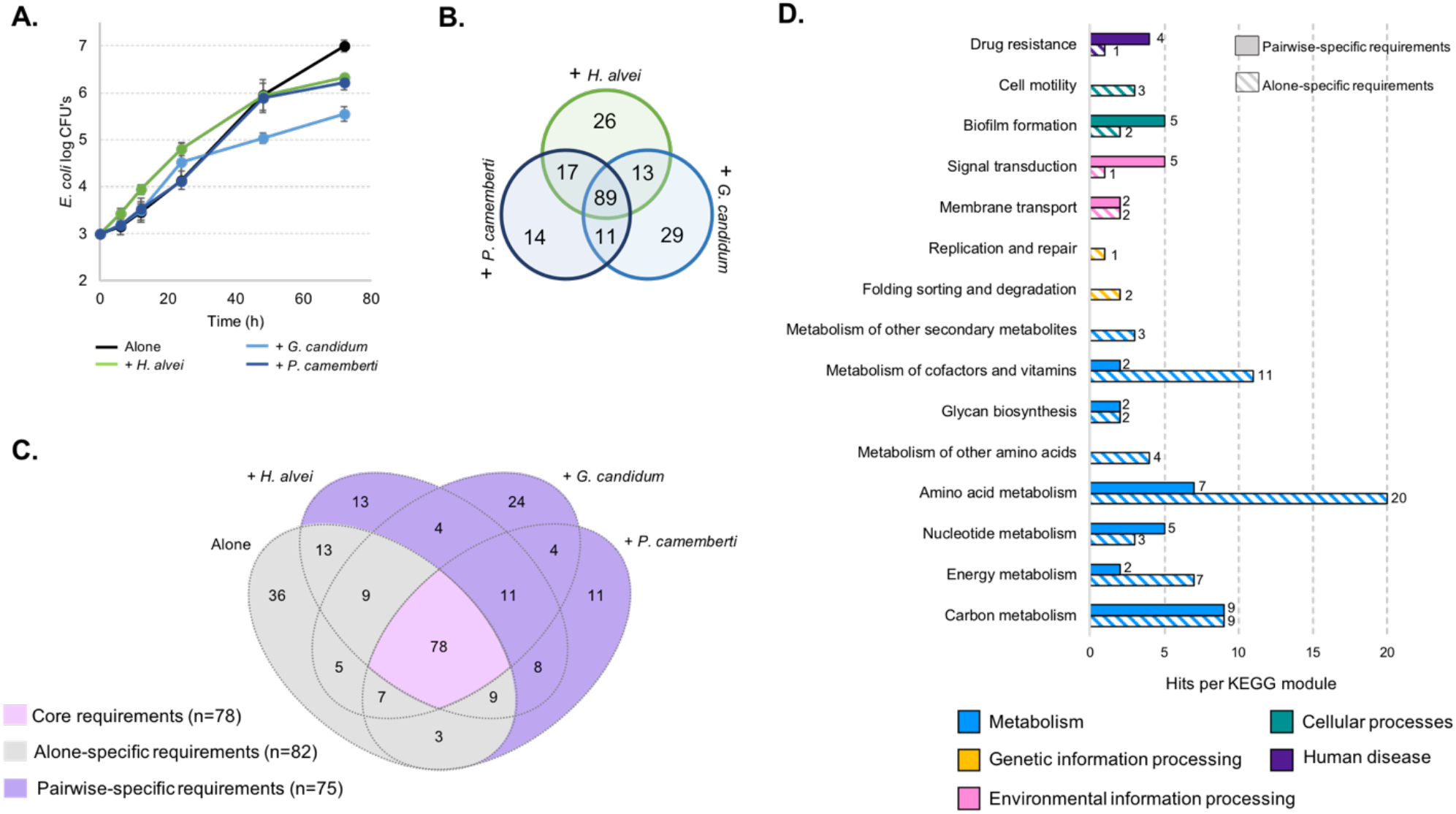
Analysis of *E. coli* essential genes during growth in pairwise conditions. We grew *E. coli* in pairwise conditions with either with *H. alvei*, *G. candidum* or *P. camemberti* **(A)**. Using the *E. coli* RB-TnSeq library we identified the essential genes for growth on CCA in each pairwise condition. Requirements for growth with the different partners overlapped between the conditions. Altogether, it constitutes 153 essential genes **(B)**. Comparing these genes to the genes essential for *E. coli* growth alone, we identified 82 genetic requirements that were no longer required for growth in pairwise conditions as well as 75 genes that were added by pairwise growth **(C)**. We mapped the pairwise-specific and alone-specific genes to the KEGG BRITE database. 45 out of 82 genes and 33 out of 77 had hits **(D).** Most of the alone-specific requirements were mapped to metabolic pathways and especially amino-acid metabolism (n=20 genes) while pairwise-specific requirements mapped to metabolic functions but also to functions associated with response to stress (biofilm formation n=5, signal transduction n=5 and drug resistance n=4).

We then determined essential genes for *E. coli* growth in each pairwise condition using RB-TnSeq (i.e any genes whose fitness value is negative and associated with an absolute t-score greater than 3 in the pairwise condition). As performed above, barcode frequencies were compared between T0 and after growth with each partner (at day 1,2 and 3). For each pairwise condition, we pooled genes with a consistent fitness for at least one timepoint as a single set of essential genes. We identified 145 conditionally essential genes for *E. coli* growth with *H. alvei*, 131 conditionally essential genes for its growth with *G. candidum* and 142 conditionally essential genes for its growth with *P. camemberti* (Figure 2B and supplementary file 2). We saw significant overlap between these gene sets, which altogether constitute a set of 153 genes that were conditionally essential for growth in at least one pairwise culture. These genes are further referred to as genes essential for growth in pairwise conditions.

Comparison of the set of genes identified when *E. coli* is grown alone with the genes identified when *E. coli* is grown in pairwise conditions is expected to highlight differences brought about by the presence of another species. Consistent presence of multiple genes of the same pathway within a set of essential genes is likely to point out a pathway specifically essential in one condition. Thus, we can infer possible interactions based on the different essential pathways between interactive and non-interactive growth conditions.

We compared the 153 genes essential in pairwise conditions to the 160 essential genes for *E. coli* growth alone (Figure 2C). Three groups of genes arose from that comparison: (i) the core requirements: genes essential for both growth alone and in pairwise conditions (n=78), (ii) the alone-specific requirements: any gene essential for *E. coli* growth alone that was not identified as essential in the presence of at least one of the partner (n=82), and (iii) pairwise-specific requirements: any gene essential in the presence of at least one of the partners but not essential for growth alone (n=75). We further focused on the alone-specific and pairwise-specific requirements as these groups contain genes potentially related to interactions.

Alone-specific requirements can highlight genes that are essential for growth alone but no longer required due to the presence of a partner, thus suggesting interactions between *E. coli* and the partner. In this specific case, these genes can be described as requirements which are relieved by the presence of a partner. We mapped the 82 alone-specific genes (Figure 2D and supplementary file 3) to the KEGG BRITE database to identify functions and pathways that are no longer essential in the presence of a partner. 16 genes were associated with unknown or predicted proteins and did not map to any field of the database. Of the remaining genes, 45 mapped to modules of the KEGG orthology hierarchy. 7 genes were associated with genetic information processing, environmental information processing, cellular processes and human disease modules. However, most of the alone-specific genes were associated with the KEGG module metabolism, and are thus part of metabolic pathways. It is especially evident that pairwise growth leads to major changes in the need for amino acid biosynthesis. For example, 6 out of the 8 genes of valine and isoleucine biosynthetic pathways are no longer required during pairwise growth (Figure 3D and supplementary file 3). In addition, 2 genes of arginine biosynthesis, 2 genes of methionine biosynthesis as well as final steps of homoserine, aspartate and glutamate biosynthesis are no longer required. Moreover, *ilvY*, the transcriptional activator of valine and isoleucine biosynthesis was also among the genes no longer required for pairwise growth. Here, the dominant presence of amino acid biosynthesis genes in the alone-specific requirements suggests cross-feeding of the pathway end-product or key intermediaries which are either provided directly by the partner species or made more available in the environment as a consequence of the partner’s metabolic activity. Thus, our data suggest that pairwise growth may allow cross feeding of the amino acids valine, isoleucine, arginine, methionine, homoserine, aspartate and glutamate. Isoleucine or methionine are also intermediaries of cofactor biosynthesis, and genes associated with their biosynthesis were also mapped to metabolism of cofactors and vitamins.

**Figure 3:**
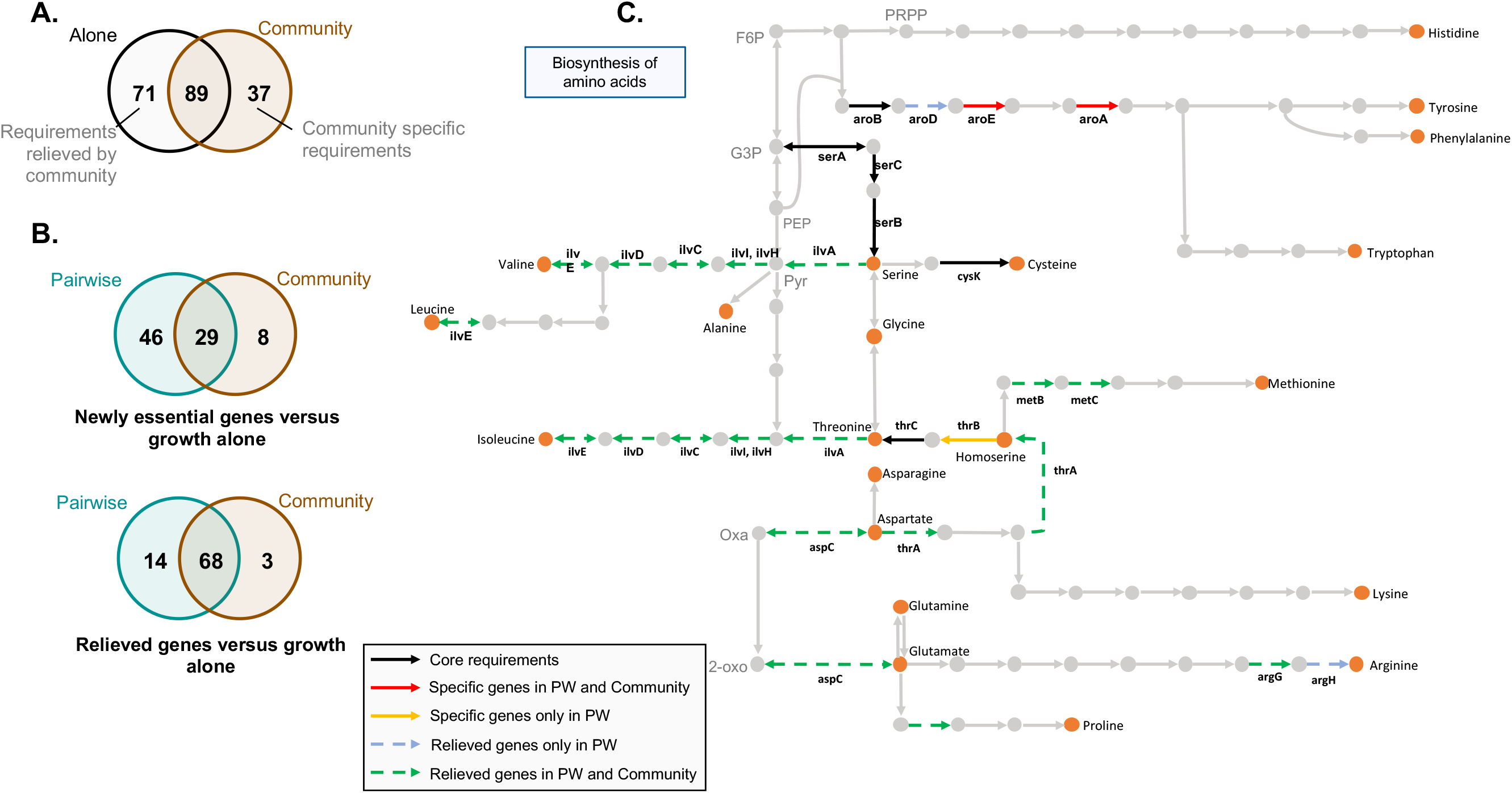
Comparison of genes essential for *E. co*li growth within the community and in pairwise conditions. Using the *E. coli* RB-TnSeq library, we identified genes required to grow with the community (*H. alvei* + *G. candidum* + *P. camemberti*). Then, we compared the essential genes for growth alone and with the community **(A)** to identify the newly essential and relieved genes during growth with the community. Next, we compared the newly essential genes in community growth versus growth alone with the newly essential genes in pairwise growth (PW) versus growth alone. We did the same for relieved requirements **(B)**. Within the requirements that were relieved both in pairwise and community growth, genes associated with numerous amino acid biosynthetic pathways were identified **(C)**. F6P: fructose-6-phosphate, PRPP: 5-phospho-ribose-1-di-phosphate, G3P: Glyceraldehyde-3-phosphate, PEP: phosphoenol-pyruvate, Pyr: Pyruvate, Oxa: Oxaloacetate, 2-oxo: 2-oxoglutarate

To understand if the alone-specific requirements were related to a specific partner, we investigated how each partner contributed to this gene set (supplementary file 3). Of the 82 total genes, 36 were identified as non-essential in all pairwise cultures. They included genes associated with amino acid metabolism specific to homoserine and methionine biosynthesis. Of the remaining genes, 8 were specifically not required in the presence of *H. alvei*, 9 were specifically not required in the presence of *G. candidum* and 9 were specifically not required in the presence of *P. camemberti*. Alleviation of leucine and valine biosynthesis was observed with all fungal partners, while biosynthesis of arginine appeared to be no longer required specifically in the presence of *G. candidum*. Fungal species are known to secrete proteases that digest small peptides and proteins (Kastman et al. 2016; Boutrou, Aziza, and Amrane 2006; Boutrou, Kerriou, and Gassi 2006) and may lead to increased availability of amino acids in the environment.

We then analyzed the 74 pairwise-specific genes (Figure 2D and supplementary file 3) in order to identify functions or pathways that are newly essential due to the presence of a partner compared. 33 genes mapped to KEGG Orthology terms. Among this gene set are pathways associated with signal transduction, biofilm formation, and drug resistance. They were related to 3 major responses: metabolic switch (*creB*: carbon source responsive response regulator), response to stress and toxic compounds (*cpxA*: sensory histidine kinase, *oxyR*: oxidative stress regulator, *acrAB*: multidrug efflux) and biofilm formation (*rcsC* and *rcsB*: regulator of capsular synthesis, *pgaC*: poly-*N*-acetyl-D-glucosamine synthase subunit). Biofilms are microbial structures known to provide resistance to different stresses, including resistance to antibiotics, and biofilm-inducing genes can be activated in presence of stress events (Landini 2009). The transcriptional regulator *oxyR* and the transduction system *CpxA* and *CpxB* are known coordinators of stress response and biofilm formation (Gambino and Cappitelli 2016; Dorel, Lejeune, and Rodrigue 2006). While these genes represent only a small subset of the all the pairwise specific gene set, they could suggest that partner species are producing toxic compounds or oxidative stress-inducing compounds.

We again investigated if these responses were partner-specific (supplementary file 3). Of the 74 pairwise-specific genes, 11 were found to be essential in the presence of all partners, 13 were specific to the presence of *H. alvei*, 24 were specific to the presence of *G. candidum* and 11 were specific to the presence of *P. camemberti*. Despite involving different genes, necessity of biofilm formation and response to toxic stress were associated with the presence of all partners.

### Identification of *E. coli* genes essential for growth within the community and comparison with genes essential for growth in pairwise conditions

We next aimed to investigate the differences between genes essential for growth in a community (complex interactive condition) and genes essential for growth in associated pairwise conditions (simple interactive condition). We grew the *E. coli* library with the complete community composed of *H. alvei*, *G. candidum* and *P. camemberti* and identified 126 essential genes (supplementary file 2). *E. coli* final biomass was reduced by the presence of the community even more so than by a single partner. However, the growth of each community member was unaffected (Supplementary figure 3).

We first compared the genes essential for growth in the community with the genes essential for growth alone. We identified 89 genes that were essential for both conditions, 37 genes only essential with the community and 71 genes that were no longer essential to grow with the community (essential only for growth alone) (Figure 3A). As for the presence of a single partner, the presence of a complex community potentially introduces new essential genes for growth while relieving some requirements.

By comparing the genes required for growth in interactive and non-interactive conditions, we have identified the set of genes which are newly required during growth in a community versus growth alone and the set of genes newly required for growth in pairwise conditions versus growth alone. Comparing these two sets of additional genes can reveal if and how community complexity modifies the genes that are essential in different interactives contexts compared to growth alone. We identified 29 essential genes that were potentially newly essential compared to growth alone in both pairwise conditions and community growth (Figure 3B and supplementary file 4). These new requirements are likely to be associated with pairwise interactions which are maintained in a community context. These include genes associated with oxidative stress and biofilm formation.

Meanwhile, we identified 46 genes that were essential only for growth in pairwise conditions, but not with the community. These genes could be related to interactions that are either alleviated or counteracted in a community, either by the presence of a specific species, or the community as a whole. For example, some of the identified genes were associated with antimicrobial resistance, and in a diverse community, other species could degrade the putative antimicrobial molecule or prevent the producing species from secreting it. Consequently *E. coli* would be exposed to a lesser level of antimicrobial, suppressing the necessity of a resistance gene.

Finally, 8 essential genes appeared to be specifically required in the presence of the community. These genes may be associated with specific interactions emerging from the community context. 3 genes encode for uncharacterized proteins, and the others (*purK*, *purE*, *damX*, *ftsX*, *secB*) are not associated with a single function or pathway. We therefore cannot conclude whether specific interactions emerge from the community context or if these genes appeared as specific because of the filtering process criteria.

We next investigated if the interactions related to relieved requirements in community growth versus alone and relieved requirements in pairwise conditions versus growth alone were similar. Thus, we compared the alone specific requirements versus growth with the community and the alone specific requirements versus growth in pairwise conditions (Figure 3B and supplementary file 4). 68 genes were no longer required for both growth in pairwise conditions and with the community compared to growth alone. These genes can also represent pairwise interactions maintained in the community context. Amino acid biosynthesis was highly represented within these genes and more specifically biosynthesis of valine, isoleucine, methionine, homoserine, aspartate and glutamate (Figure 3C). This suggests that, despite the presence of more species, these amino acids are still cross-fed.

We also identified 14 genes that were no longer required in pairwise conditions compared to growth alone yet remained essential for growth with the community. Finally, 3 genes appeared to be specifically no longer required in the presence of the community. In both cases, too few genes are involved for us to infer any hypothesis on the existence of specific pairwise interactions not conserved in the community or the emergence of specific interactions in the community.

To summarize, the genes that would be newly essential in the community compared to growth alone were mostly maintained from newly essential genes identified in pairwise conditions compared to growth alone. Similarly, the genes that were no longer essential for the growth in the community were highly similar to the genes that were no longer essential in pairwise condition. However, many newly essential genes for growth in pairwise condition compared to growth alone were not found as essential for growth within the community. Altogether, this highlights that part of the pairwise interactions are conserved within a community while underlining the presence of higher order interactions related to the higher complexity of the growth condition.

### E. coli differential expression analysis in pairwise conditions versus growth alone

So far, we used a genome-scale genetic approach to investigate potential microbial interactions. As a complementary strategy, we generated transcriptomic data for *E. coli* during growth in each previously discussed condition. Changes in transcriptional profiles can be a powerful indicator of an organism’s response to an environment and were previously used to identify *E. coli* pathways involved in interactions (Croucher and Thomson 2010; McAdam, Richardson, and Fitzgerald 2014; Galia et al. 2017).

To measure *E. coli* gene expression, we extracted and sequenced RNA from each timepoint and condition of the samples used for RB-TNseq above (after 1, 2 and 3 days of growth when grown alone, in pairwise conditions or with the community). Comparison of transcriptional profiles suggests a strong reorganization of *E. coli* gene expression over time and in response to the presence of a partner. Principal component analysis on the normalized gene expression values (rlog transformation(Love et al. 2015)) revealed that samples clustered based on the growth timepoint (separation on the horizontal axis) and whether *E. coli* is growing alone or not (separation on the vertical axis) (Figure 4A). The different pairwise conditions and the community conditions cluster together, underlining that the potential differences between these conditions are less important than the differences between an alone and co-cultured condition.

**Figure 4:**
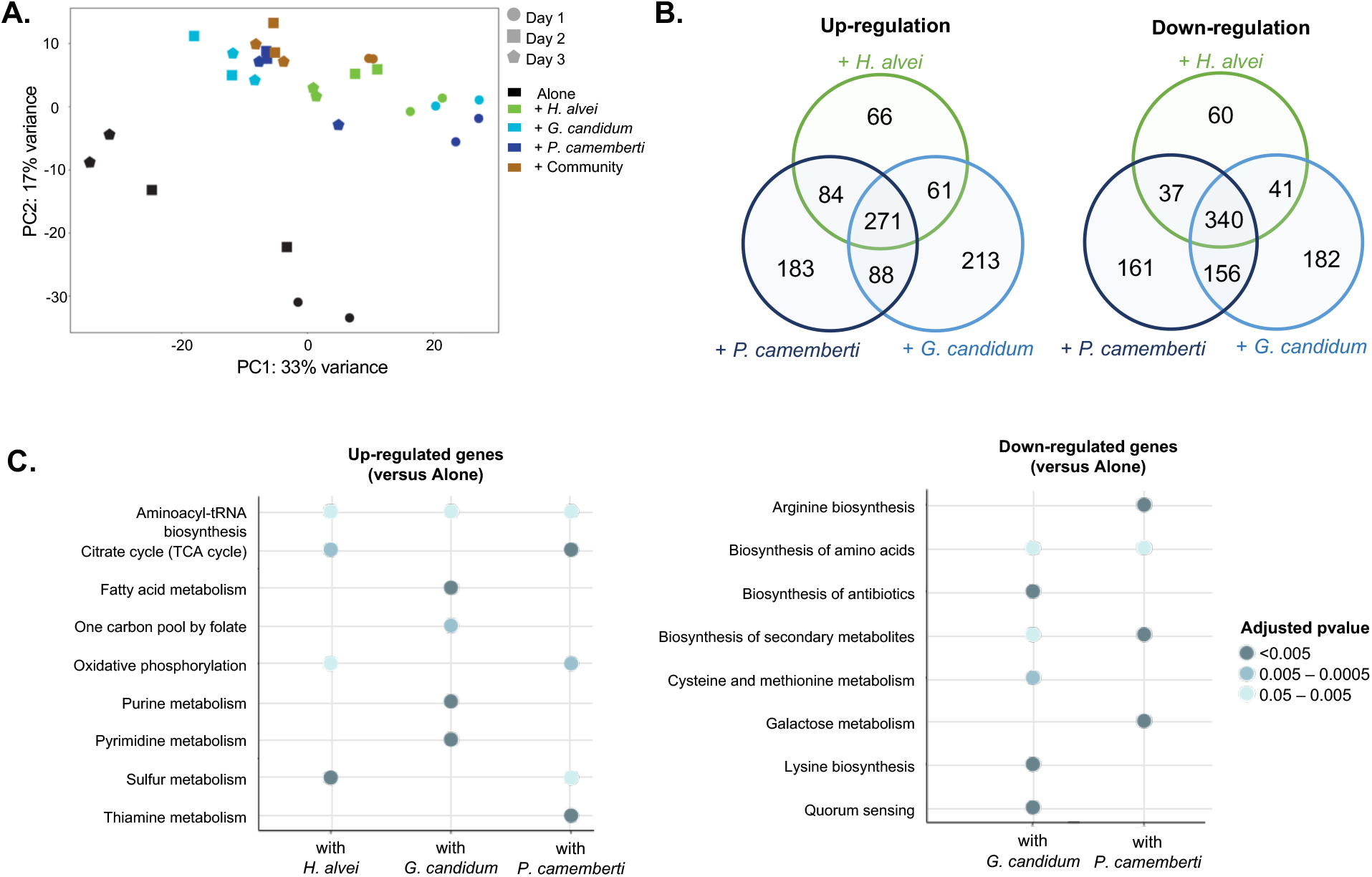
*E. coli* gene expression analysis during growth on CCA alone, in pairwise conditions and with the community. We used RNASeq to investigate *E. coli* gene expression at three timepoints (1, 2 and 3 days) during growth on CCA alone, in pairwise conditions (with *H. alvei*, *G. candidum* or *P. camemberti*) and with the community. We carried out a principal component analysis on the rlog transformed expression data (Love et al., 2015) **(A)**. Using DESeq2 (Love et al., 2015), we identified up and down-regulated genes in each pairwise condition at each timepoint compared to growth alone. We kept only genes associated with a p-value lower than 1% and an absolute logFC higher than 1. For each pairwise condition, we pooled together up-regulated genes at any timepoint and did the same for down-regulated genes. Comparing the up-or down-regulated genes in the different pairwise conditions, we highlighted overlapping response as well as specific responses depending on the partner **(B)**. Then, we carried out functional enrichment analysis on the KEGG pathways for up-or down-regulated genes in each pairwise condition. To do so, we used the R package clusterProfiler (Yu et al., 2012) as well as Benjamini-Hochberg multiple comparison correction. Only the Kegg pathways enriched with a p-value lower than 5% were considered **(C)**.

We next focused on the genes differentially expressed between pairwise and alone growth and by calculating the fold change of gene expression between pairwise growth and growth alone. We identified differentially expressed genes by screening for adjusted p-values lower than 1% and absolute log of fold change (logFC) greater than 1. To remain consistent with the analysis performed for the genetic requirements, we pooled the data at all time points after identifying all of the up-regulated or down-regulated genes for each timepoint. We found a total of 966 upregulated and 977 downregulated across all partners (482 upregulated genes and 478 downregulated genes in presence of *H. alvei*, 633 upregulated genes and 719 downregulated genes in presence of *G. candidum*, 626 upregulated genes and 694 downregulated genes in presence of *P. camemberti*, Figure 4B and supplementary file 5). 271 genes were up-regulated in all pairwise conditions while 340 genes were always down-regulated. Meanwhile, a number of genes were differentially expressed depending on the partner: 66 genes were specifically up-regulated and 60 genes down-regulated with *H. alvei*, 213 up-regulated and 182 down-regulated with *G. candidum*, 183 up-regulated and 161 down-regulated with *P. camemberti*.

Due to the larger gene set compared to RB-TNseq, we performed KEGG Pathway enrichment analyses on the differentially expressed genes in pairwise conditions to determine up-regulated functions and pathways depending on the partner species (Figure 4C). First, certain functions are up-regulated regardless of the partner. This includes up-regulation of almost all the aminoacyl-tRNA-synthetases and up-regulation of energy production. Interestingly up-regulation of energy production through aerobic respiration and the TCA cycle happened after 3 days of growth. Oxygen availability (Gunsalus 1992) and growth phase (Wackwitz et al. 1999) are the two known regulators of aerobic respiration. At day 3, *E. coli* was observed to be in log phase when alone, whereas in the presence of a partner and especially with *P. camemberti*, *E.coli* was observed to enter the stationary phase between day 2 and day 3 (Supplementary figure 5). Therefore, up-regulation of aerobic respiration is most likely associated with the growth stage difference between *E. coli* alone and with a partner.

More genes were up-regulated in the presence of *G. candidum* than the other partners. Several pathways associated with nucleotide biosynthesis (one-carbon pool by folate, purine metabolism and pyrimidine metabolism) were specifically upregulated. This suggests that either *coli* and *G. candidum* compete for nucleotide compounds from the environment or that presence of *G. candidum* leads to an increase demand of nucleotide compounds for *E. coli’s* metabolism and growth.

We performed a similar KEGG pathway enrichment analysis on the down-regulated genes in pairwise conditions. No pathway was enriched in the presence of *H. alvei.* Pathways involved in the biosynthesis of amino acids, specifically tyrosine, phenylalanine, tryptophan, methionine, lysine, arginine homoserine, leucine, glutamate, threonine and glycine, appeared to be the principal down-regulated function in the presence of a fungal partner. Lysine biosynthesis was specifically down-regulated with *G. candium* while arginine biosynthesis was specifically downregulated with *P. camemberti*. Interestingly, some amino acid biosynthetic pathways were up-regulated later in the growth (phenylalanine, tyrosine and leucine). Down-regulation of amino acid biosynthesis suggests that the partner species generates amino acid cross feeding. Our previous RB-TnSeq results previously suggested this particular interaction in the presence of fungi. The observation of this interaction in the transcriptome data is consistent with our interpretation of RB-TnSeq and reinforces the likelihood of such an interaction. However, late up-regulation of some amino acid biosynthesis suggests that as the partner grows along with *E. coli* they eventually end up competing for amino acids leading to biosynthesis up-regulation.

To summarize, presence of a single partner triggers a deep and dynamic reorganization of *E. coli* gene expression. These modifications mostly rely on restructuring *E. coli* metabolic activity whether it is to down-regulate specific pathways and activate new ones as a response to potential metabolic competition and establishment of an appropriate metabolic strategy or to down-regulate certain pathways and benefit from cross-feeding and common goods.

### Comparison of differentially expressed genes in pairwise condition versus alone and community growth versus alone

To determine whether *E. coli* gene expression significantly changes when grown with the full community as compared to growth in pairwise conditions, we first calculated *E. coli* gene logFC at each timepoint between growth with the community and growth alone. We further analyzed genes with adjusted p-values lower than 1% and absolute logFC greater than 1. After pooling across timepoints, we identified 465 up-regulated and 476 down-regulated genes in the presence of the community versus growth alone (Supplementary file 5). We then compared these genes to the 966 up-regulated genes and 977 down-regulated genes in pairwise conditions versus growth alone (Figure 5A).

**Figure 5:**
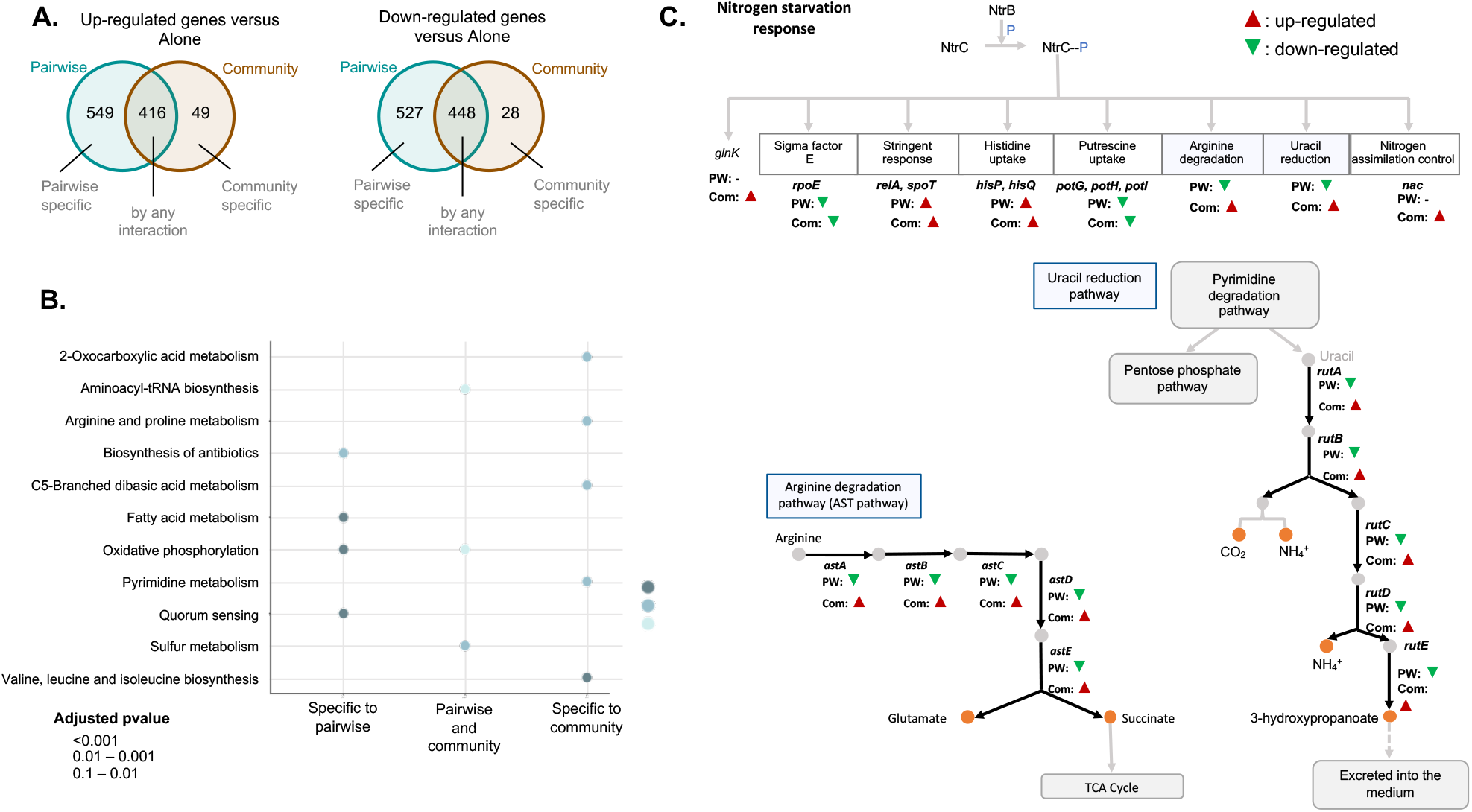
Comparison of differentially expressed genes in pairwise conditions versus alone and community growth versus alone. Using DESeq2 (Love et al., 2015), we identified up and down-regulated genes during growth with the community compared to growth alone at each timepoint. We kept only genes associated with a p-value lower than 1% and an absolute logFC higher than 1. We pooled together up-regulated genes at any timepoint and did the same for down-regulated genes. Then, we compared up-regulated genes in pairwise conditions with genes up-regulated during growth with the community and down-regulated genes in pairwise conditions with down-regulated genes during growth with the community **(A)**. We further performed functional enrichment analysis on KEGG pathways using the R package clusterProfiler (Yu et al., 2012) on the genes differentially expressed in different situations. We used Benjamini-Hochberg multiple comparison correction and only the Kegg pathways enriched with a p-value lower than 5% were considered **(B)**. Within the genes specifically up-regulated during growth with the community, we observed the up-regulation of multiple genes associated with the nitrogen starvation response **(C)**.

First, we identified 549 genes that were specifically up-regulated in pairwise conditions versus growth alone and not up-regulated in community versus growth alone. KEGG pathway enrichment analysis highlighted that these genes were mostly associated with quorum sensing, fatty acid metabolism and oxidative phosphorylation (Figure 5B). This observation suggests that addition of other species in the community counteracts or prevents pairwise specific interactions from that partner. 416 genes were found to be up-regulated both in pairwise and community growth versus growth alone. Enrichment analysis highlighted functions that were previously described as up-regulated in most of the pairwise conditions: aminoacyl-tRNA-synthetase and energy metabolism (Figure 5B). This suggests that certain interactions that *E. coli* experienced in pairwise conditions are conserved in the community context. To investigate if the addition of potentially similar interactions from different partners leads to an amplified response, we explored if the magnitude of expression changes in these pathways is higher in the community. We performed differential expression analysis on the genes comparably regulated in pairwise conditions and with the community (Supplementary file 6). 50 of the 416 up-regulated were significantly more up-regulated in community growth compared to pairwise growth. Among them, sulfate assimilation genes were over-represented. This suggests that similar pairwise interactions may be additive in the community leading to a stronger transcriptional response.

Finally, 49 genes were specifically up-regulated with community growth versus growth alone. Despite a low number of genes, KEGG pathways were significantly enriched (Figure 5B). These enriched groups include the biosynthesis of valine, leucine and isoleucine, pyrimidine metabolism as well as arginine and proline metabolism. Up-regulation of certain amino acid biosynthesis pathways suggests that despite potential cross feeding from individual partners, addition of many partners eventually leads to competition. Up-regulation of pyrimidine, arginine and proline metabolism however is part of a larger response; the response to nitrogen starvation (Figure 5C). This response facilitates cell survival under nitrogen-limited conditions. Specifically, up-regulated genes included all the genes involved in the regulatory loop of the transcriptional regulator NtrC (*glnL*) and nitrogen utilization as well as NtrC transcriptional targets: the transcriptional regulator Nac (*nac*), the operon *rutABCDEFG* involved in ammonium production by uracil catabolism, the *astABCDE* operon constituting the arginine degradation pathway (AST pathway) and the two regulators of the stringent response, *relA* and *spoT*. Thus, addition of species in the community specifically triggers the activation of the response to nitrogen starvation, which suggests a potential higher competition for nitrogen in the community context.

We performed a similar analysis on down-regulated genes in pairwise conditions and with the community versus growth alone to investigate if transcriptional down-regulation in pairwise and community conditions are similar (Figure 5A). 527 genes appeared specifically down-regulated in pairwise conditions versus growth alone. Despite the large number of genes, no KEGG pathway was significantly enriched among these genes. However, the *rutABCDEFG* and *astABCDE* operons associated with the response to nitrogen starvation were down-regulated in each pairwise conditions (Figure 5C). This observation strongly suggests that presence of the community triggers a highly specific response that would be down-regulated in the presence of only one species.

We identified 448 genes that were down-regulated during both pairwise and community growth conditions versus growth alone. Enrichment analysis pointed to the down-regulation of amino acid biosynthesis, lysine biosynthesis as well as cysteine and methionine metabolism. Therefore, consistent with our RB-TnSeq data, this suggests that cross-feeding from a single partner is maintained in a more complex context. Finally, 28 genes were specifically down-regulated when *E. coli* was grown with the community. Individual analysis of these genes highlighted 9 genes that code for uncharacterized proteins but did not highlight over-represented functions.

To conclude, most of the changes in *E. coli* gene expression during growth with the community were similar to those observed in pairwise conditions. Moreover, some of these changes were amplified in the community compared to pairwise. This suggests that while a large part of transcriptional regulation in community results from pairwise interactions, similar interactions from different partners may be additive in the community and exert a stronger impact on transcription. Also, the observed changes in nitrogen availability-related transcription suggest that community growth may induce new metabolic limitations.

### RB-TnSeq identification of genes essential for *Pseudomonas psychrophila* JB418 growth with the bloomy rind cheese community

Because *E. coli* is a well-characterized model organism, it provides a good starting point for genetic analysis of interactions in this system. However, because *E. coli* is not typically found in cheese, we wanted to characterize interactions using an endogenous species. We selected *Pseudomonas psychrophila* JB418, which was previously isolated from a bloomy rind cheese sample (Wolfe et al. 2014), and is genetically tractable,

We first sequenced, de novo assembled, and annotated the genome of this species (see Materials and Methods). We used KEGG KOALA Blast (Kanehisa, Sato, and Morishima 2016) to obtain KEGG identifiers for coding sequences. 2983 coding sequences (51.5%) were assigned a KEGG identifier. We then used the same barcoded, transposon delivery library to construct a RB-TnSeq library in this strain (see materials and methods for library construction and characterization). We used TnSeq analysis to map barcode insertions to the annotated genome. 272,329 insertions were mapped and 143,491 were centrally inserted (transposons mapping to the first and last 10% of a gene were discarded) at 77,766 different locations. With 4811 genes with at least one central insertion, the library covers 83% of *P. psychrophila* genome (Figure 6A). The genes with no central insertion are likely to be essential genes for growth on LB.

**Figure 6:**
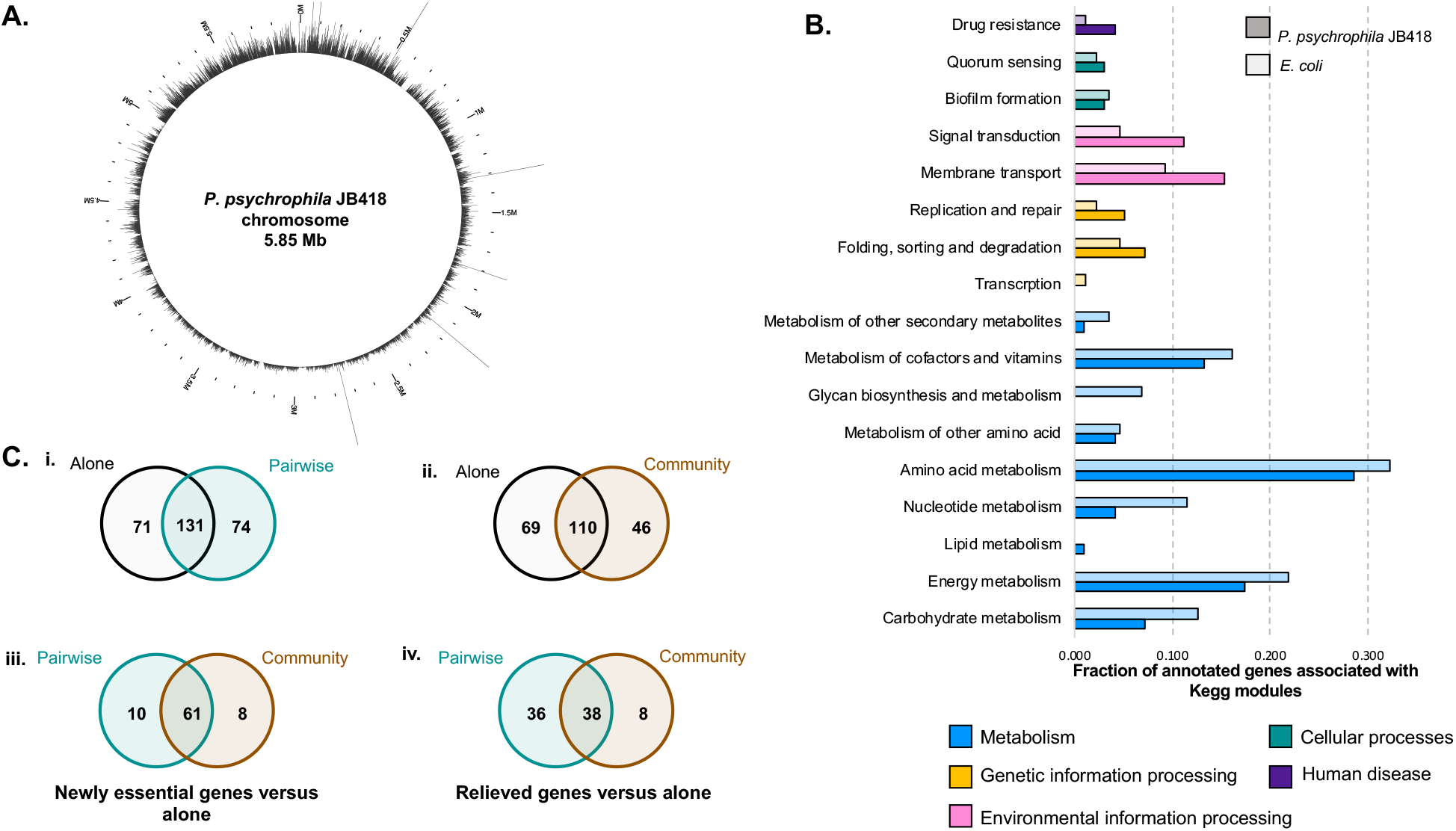
Identification of genes essential for *P. psychrophila* JB418 growth on CCA and with the model community. We built a barcoded-transposon library in the cheese isolate *P. psychrophila* JB418. 272,329 insertions were mapped to the genome. The library covers 83% of *P. psychrophila* JB418 genome. On the chromosome, each bar represents the number of insertions per kB **(A)**. We grew the library alone on CCA and identified 179 genes essential for *P. psychrofila* JB418. We annotated *P. psychrophila* JB418 using KEGG KOALA Blast (Kanehisa, Sato, and Morishima 2016). 98 of the 179 genes were attributed a KEGG annotation. To compare the relative importance of essential functions for growth alone observed for *P. psychrophila* JB418 and for *E. coli* we represented, per KEGG module, the ratio between the number of hits for the module and the number of genes with KEGG annotation **(B)**. We further identified *P. psychrophila JB418* essential genes for growth in the different pairwise conditions and with the community. Then, we determined which genes were newly essential or relieved in pairwise conditions compared to growth alone as well as newly essential or relieved genes with the community compared to growth alone. Finally, we compared newly essential genes in pairwise conditions versus alone with newly essential genes in the community versus alone as well as relieved genes in pairwise condition versus alone with relieved genes in community versus alone **(C)**.

We then grew the *P. psychrophila* library for 3 days on CCA either alone (supplementary figure 4), in pairwise conditions (with *H. alvei*, *G. candidum* or *P. camemberti*) and with the full community. Fitness values were successfully calculated for 3060 genes and the same statistical screening (absolute t-value >=3) and fitness screening (fitness value < 0) were used to identify essential genes. We first analyzed the 179 genes essential for *P. psychrophila* growth alone (Figure 6B and supplementary file 7). Most of the essential genes for *P. psychrophila* growth alone on CCA appeared to be associated with metabolism. More specifically, they are predicted to be involved in amino acid biosynthesis (all amino acids except for lysine), biosynthesis of cofactors and vitamins as well as energy metabolism (sulfate assimilation). Also, pathways associated with membrane transport were related to iron-siderophore uptake and glycine betaine transport. Essential functions for *P. psychrophila* growth alone were consistent with the previous functions identified for *E. coli* growth alone.

Next, we identified 152 genes essential for *P. psychrophila* JB418 growth with *H. alvei*, 176 genes essential for growth with *G. candidum*, and 164 genes essential for growth with *P. camemberti* (Supplementary file 7). Altogether, they represent 205 genes essential for *P. psychrophila* growth in pairwise conditions. Compared to growth alone, 71 genes are no longer required for *P. psychrophila* during pairwise growth (Figure 6Ci). We mapped these relieved genes to the KEGG BRITE database to identify potentially relieved functions. Of the 71 genes, only 27 were assigned a KEGG annotation. Here, no pathway appeared to be clearly relieved (Supplementary figure 5).

We identified 74 newly essential genes for *P. psychrophila* growth in pairwise conditions compared to growth alone (Figure 6Ci) and 34 of them were mapped to the KEGG Brite DataBase (Supplementary figure 5). These genes included 11 associated with aromatic amino acid biosynthesis. Since amino acid biosynthesis is also required for growth alone, these data suggest that introduction of a partner leads to an amplified depletion of free amino acids, likely due to competition for these nutrients. Additionally, 4 genes associated with DNA repair were identified as essential in the presence of another species. This could suggest presence of a toxic stress produced from the partner.

Finally, we identified 156 genes essential for *P. psychrophila* when grown with the community (Supplementary file 7). Compared to growth alone, we identified 69 genes that were no longer required (Figure 6Cii). Then, comparing these 69 relieved genes in community versus growth alone with the relieved requirements in pairwise conditions versus growth alone (Figure 6Ciii), we concluded that only 8 genes were no longer required uniquely in the community. 4 of these 8 genes were associated with valine and isoleucine biosynthesis, suggesting that addition of more species might lead to cross-feeding of these amino acids similar to what was observed with *E. coli*.

We identified 46 genes that were newly essential for *P. psychrophila* growth with the community versus growth alone (Figure6Cii). Comparing these genes to the newly essential genes identified in pairwise growth versus growth alone, we identified 8 genes uniquely essential during community growth and not during any pairwise growth condition (Figure 6Civ). Of these genes, 4 were annotated as either transporters or putative transporters. Here, as for *E. coli*, we observed that newly essential and relieved requirements in the community context were mostly maintained from pairwise conditions, while not all pairwise newly essential or relieved requirements were maintained in the community.

To summarize, essential functions for *P. psychrophila* growth alone on CCA were consistent with the ones identified for *E. coli*. While demonstrating important environmental parameters to be considered for growth on CCA such as low available iron, low available free amino acid and high osmolarity, it also underlines the robustness of the RB-TnSeq approach in accurately identifying essential genes. Analysis of the functions of newly essential or relieved genes in interactive contexts allowed us to examine potential *P. psychrophila* interactions with the members of our model community. Interestingly, the possible interactions probed with *P. psychrophila*, whether they are associated with relieved essential genes or newly essential genes, appeared to differ from the ones identified with *E. coli*.

## DISCUSSION

In this work, we combined high-throughput, genome scale approaches to begin to define microbial interactions within a model community. Our integrative approach relies on the use of the model species *E. coli* as a genetic probe grown either alone, in pairwise culture with each community member or with the complete community. We first used RB-TnSeq to determine *E. coli’s* genetic requirements in these different contexts associated with increasing levels of complexity. Our analysis showed that 3.86% of the genes in the *E. coli* genome were important to grow in our experimental environment. 53.1% of these genes appeared to be no longer required when grown with a partner but a distinct set of genes (comprising 1.98% of the *E. coli* genome) appeared to be specifically required during interactive conditions. As identification of differences for genes essential across conditions is based on a stringent filtering cutoff for consistent fitness values, some differences might be attributed to false negative results. For example, during growth alone, 11 genes that were found to be essential only at day 1 become essential at all timepoints if we relax our t-score cutoff to absolute t-score >= 2 and increase our false discovery rate to 2%. However, essentiality differences between timepoints can also point out that genetic requirements do change over time. A method for quantitative comparison of fitness values across conditions or timepoints will need to be developed to overcome that issue. However, since we expect the false negative results to be randomly distributed across *E. coli*’s genes, we consider that if genes belonging to the same pathway or function are specially observed in a condition and not another, they are very likely to be involved in a microbial interaction. To complement the RB-TnSeq approach, we used RNASeq and differential gene expression analysis to investigate *E. coli* transcriptional response during growth in pairwise culture and with the community compared to a growth alone. Here, we highlighted a deep reorganization of gene expression whenever *E. coli* is in the presence of other species.

Our work illustrates for the first time the use and complementarity of RB-TnSeq and RNASeq for microbial interaction analysis. These techniques lead to consistent interpretations and each method compensated for the limitations of the other. RNASeq provides a comprehensive view of interactions by looking at the transcriptional response of *E. coli* to an interactive context. However, it fails to provide information on the actual requirement of a given gene or pathway for an interactions. RB-TnSeq identifies the interactions that have a strong impact on the *E. coli* growth phenotype including genes for which expression is constitutive in all conditions and won’t be highlighted with RNASeq analysis. On the other hand, gene redundancy and the potential presence of “common good” molecules in the library are expected to prevent identification of certain interactions by RB-TnSeq but not by RNASeq. This was illustrated by the absence of essentiality of the biosynthesis and secretion of enterobactin but the essentiality of enterobactin uptake system. It is very likely that the majority of the library synthesizes and secretes enterobactin and thus the enterobactin synthesis mutants can uptake this “common good” and survive. Interestingly, RNASeq analysis demonstrated upregulation of the enterobactin biosynthesis pathway, suggesting enterobactin synthesis is necessary in an interactive context. Similarly, differential expression analysis of the genes highlighted by RB-TnSeq as essential in any condition allowed us to explore the presence of interactions associated with these core functions. For example, it highlighted that iron-uptake was amplified during interactions suggesting competition for that already limiting nutrient.

Together, interpretation of the functional analysis of RB-TnSeq and RNASeq results pointed to possible microbial interaction mechanisms in the different growth contexts. Addition of genetic requirements associated with metabolism and/or up-regulation of similar functions can be interpreted as presence of metabolic competition toward associated compounds. Conversely, alleviation of biosynthetic genes and/or downregulation of biosynthetic functions can point out cross-feeding of associated compounds. Finally, production of a specific stress by the partner can be deduced from addition of requirements associated with stress response and/or up-regulation of similar functions.

Combining RB-TnSeq and RNASeq analysis, we identified potential interactions in pairwise and community conditions. We have observed competition for iron in most interactive conditions and in pairwise conditions, the partner appeared to produce possible toxic compounds. Analysis of available genomes of *H. alvei* (Tan, Yin, and Chan 2014), *G. candidum* (Polev et al. 2014) and *P. camemberti* (Cheeseman et al. 2014) in AntiSmash (Weber et al. 2015) highlighted the ability of each species to produce extracellular metabolites with potential toxic properties including bacteriocins for *H. alvei* and diverse polyketides and terpenes for the fungi. Analysis of extracellular metabolites will be necessary to characterize the molecular signature of these stresses. Finally, we consistently observed possible amino acid cross feeding from fungal partners, either in pairwise or community growth. This is very likely due to the fungal capacities of secreting proteases and digesting the small peptides and proteins present in the environment (Kastman et al. 2016; R. Boutrou, Aziza, and Amrane 2006; Rachel Boutrou, Kerriou, and Gassi 2006). The observation of cross-feeding strengthens the statement that cross-feeding events are widespread within communities (Goldford et al. 2017).

In our bottom-up approach, RNASeq and RB-TnSeq consistently highlighted that only a fraction of pairwise interactions were conserved in the community. This highlights the existence of higher order interactions counteracting simple pairwise interactions (Billick and Case 1994). Higher order interactions could be compensatory pairwise interactions or upstream effects ending in pairwise interaction cancellation. Although community structure is argued to be predictable from pairwise interactions in specific cases, higher order interactions are believed to be responsible for the general lack of predictability (Billick and Case 1994; Friedman, Higgins, and Gore 2017; Momeni, Xie, and Shou 2016). Here, our work demonstrated the existence of these higher order interactions even within a very simple community.

We generated data using both *E. coli*, a non-endogenous species, and *P. psychrophila* JB418, a cheese endogenous species. We identified similar functions in both species as important requirements to grow in the cheese environment. However, growth in interactive contexts highlighted different interactions. We hypothesize that either *E. coli* and *P. psychrophila* do not similarly cope with the presence of a given partner or the partner species response depends on the probe-species as well as a combination of both scenarios. Altogether, this suggests important species-specific interaction patterns. However, our analysis was limited, especially for *P. psychrophila*, by genes which were poorly annotated or not characterized. Further investigation and characterization of these genes would improve interaction characterization and may highlight new microbial interactions pathways.

This study represents a novel approach to investigating microbial interactions in communities, and revealed the intricacy, redundancy and specificity of the many interactions governing a simple microbial community. The ability of *E. coli* to act as a probe for molecular interactions, the robustness of RB-TnSeq, and its complementarity with RNASeq open new paths for investigating molecular interactions in more complex communities, independently of their genetic tractability, and can thus contribute to a better understanding of the complexity and diversity of interactions within microbiomes.

## MATERIAL AND METHODS

### Strains and media

The following strains have been used to reconstruct the bloomy rind cheese community: *H. alvei* JB232 isolated previously from cheese (Wolfe et al. 2014) and two industrial cheese strains: *G. candidum* (*Geotrichum candidum* GEO13 LYO 2D, Danisco – CHOOZIT^TM^, Copenhagen, Denmark) and *P. camemberti* (PC SAM 3 LYO 10D, Danisco - CHOOZIT^TM^). The strain *P. psychrophila* JB418 was isolated from a sample of Robiola due latti (Italy) previously (Wolfe et al) and used for all the experiments involving *Pseudomonas*. All the *E. coli* strains used in this study shared the same genetic background of the initial strain *E. coli* K12 BW25113. The use of the different strains is described in Table 1.

**Table 1:** E. coli strains used during the study.

| Experiment | *E. coli* strain(s) | Ref |
| --- | --- | --- |
| RB-TnSeq | *E. coli* Keio_ML9 library | (Wetmore et al. 2015) |
| Growth assays | *E. coli* “JW0024” strain (undisrupted mutant) | (Baba et al. 2006) |
| Competition assays | WT: Keio ME9062<br>Mutants: (supplementary figure 2) | (Baba et al. 2006) |

### Growths assays on 10% cheese curd agar, pH7

All growth assays have been carried out in at least triplicates on 10% cheese curd agar, pH7 (CCA) (10% freeze-dried Bayley Hazen Blue cheese curd (Jasper Hill Farm, VT), 3% NaCl, 0.5% xanthan gum and 1.7% agar). The pH of the CCA was buffered from 5.5 to 7 using 10M NaOH.

#### *E. coli* and *P. psychrophila* JB418 growth on CCA

Growth assays have been carried out for the *E. coli* “JW0024” strain (Baba et al., 2006) and *P. psychrophila* JB418. For each species, 7*10^7^ cells were inoculated on the surface of a Petri dish containing 20mL of CCA (12.7*10^4^cells/cm^2^) after an overnight pre-culture in liquid LB-kanamycin (50µg/ml) at 37°C for *E. coli* or at room temperature (RT) for *P. psychrophila* JB418. Growth on CCA was then carried out for 3 days at RT. Plugs of 0.44cm^2^ were removed from the Petri dish at T= 0h, 6h, 12h, 24h, 36h, 48h, 72h, 120h and 240h for *E. coli* and T=0h, 12h, 24h, 48h, 72h and 240h for *P. psychrophila* JB418. They were homogenized in 1 mL of PBS1X-Tween0.05% and 3 dilutions were plated on LB-kanamycin (50µg/mL) for *E. coli* or LB for *P. psychrophila*. Plates were incubated for 24h at 37°C for *E. coli* and at RT for *P. psychrophila* JB418. After incubation, colony forming units (CFU) counting was used to estimate the number of bacterial cells on the cheese curd agar plates.

#### *E. coli* growth in pairwise cultures

Pairwise cultures were carried out on 96 well plates containing 200µL of CCA per well. *E. coli* was co-inoculated with either *H. alvei*, *G. candidum* or *P. camemberti* at a ratio of 1:1 cells (1000 cells of *E. coli* and 1000 cells of the partner). Single growth of *H. alvei*, *G. candidum* and *P. camemberti* were also measured by inoculating 2000 cells of each species on individual wells. At T=0h, 6h, 12h, 24h, 36h, 48h and 72h, a pellet of each condition was harvested, homogenized in 1 mL of PBS1X-Tween0.05% and 3 dilutions were plated on LB, LB-kanamycin (50µg/mL), LB-chloramphenicol (50µg/mL) and/or LB-kanamycin:cyclohexamide (50µg/mL and 10µg/mL) to follow the growth of each species (see Table 2). Plates were then either incubated overnight at 37°C (LB, LB-kanamycin and LB-kanamycin:cyclohexamide plates) or for 2 days at RT (LB-chloramphenicol plates) before determination of CFUs.

**Table 2:** Organization of CFU’s quantification for growth assays.

| Growth assay | Plates used |
| --- | --- |
| *E. coli* + *H. alvei* JB232 | LB (*E. coli* + *H. alvei* JB232 CFU's) LB-kanamycin (*E. coli* CFU's) |
| *E. coli* + *G. candidum* | LB-kanamycin:cyclohexamide (*E. coli* CFU's) LB-chloramphenicol (*G. candidum* CFU's) |
| *E. coli* + *P. camemberti* | LB-kanamycin:cyclohexamide (*E. coli* CFU's) LB-chloramphenicol (*P. camemberti* CFU's) |
| *H. alvei* | LB |
| *G. candidum* | LB-chloramphenicol |
| *P. camemberti* | LB-chloramphenicol |
| *E. coli* + Community | LB-cyclohexamide (*E. coli* and *H. alvei* JB232 CFU's), LB-kanamycin:cyclohexamide (*E. coli* CFU's), LB-chloramphenicol (*G. candidum* and *P. camemberti* CFU's) |

#### *E. coli* growth in the bloomy rind cheese community

Cultures were grown on 96 well plates containing 10% cheese curd agar at pH7. The four species (*E. coli*, *H. alvei* JB232, *G. candidum* and *P. camemberti*) have been co-inoculated at a ratio of 10:10:10:1. At T=0h, 6h, 12h, 24h, 36h, 48h and 72h, a pellet was harvested, homogenized in 1 mL of PBS1X-Tween0.05%, serially diluted and plated on LB-cyclohexamide (10µg/mL), LB-chloramphenicol (50µg/mL) and LB-kanamycin:cyclohexamide (50µg/mL and 10µg/mL) (see Table 2) to follow the growth of each species. Plates were then either incubated overnight at 37°C (LB-cyclohexamide and LB-kanamycin:cyclohexamide plates) or for 2 days at RT (LB-chloramphenicol plates) before CFU counting. *P. camemberti* and *G. candidum* colonies are morphologically distinct and CFUs of each species could be counted from the same plate.

### *P. psychrophila* JB418 genome sequencing, assembly and annotation

*P. psychrophila* JB418 gDNA was sequenced using Pacific Biosciences (PacBio), Oxford Nanopore Minion (Oxford Nanopore, Oxford, UK), and Illumina sequencing. PacBio library preparation and sequencing were performed by the IGM Genomics Center at the University of California, San Diego. Nanopore library preparation and sequencing were done at the University of California, Santa Barbara as part of the KITP Quantitative Biology summer course. Illumina library preparation and sequencing were done at the Harvard University Center for Systems Biology. Canu was used to assemble the PacBio and nanopore reads (Koren et al. 2017). Pilon was then used to polish the assembly using Illumina genomic sequencing data (Walker et al. 2014). The assembled genome was annotated using the Integrated Microbial Genomes & Microbiomes (IMG/M) system (Markowitz et al. 2012). The *P. psychrophila* JB418 genome is 6,072,477 nucleotides long. It contains a single circular chromosome of 5.85 Mb and 4 plasmids of 172.2 Kb, 37.7Kb, 5.8Kb and 2.4Kb. 6060 genes including 5788 open reading frames were identified.

### Transposon mutant library construction in *P. psychrophila* JB418

*P. psychrophila* JB418 was mutagenized by conjugation with *E. coli* strain APA766 (donor WM3064 which carries the pKMW7 Tn5 vector library) (Wetmore et al. 2015). This donor strain is auxotrophic for diaminopimelic acid (DAP). The full collection of the APA766 donor strain (1mL) was grown up at 37°C overnight at 200 rpm. Four 25mL cultures (each started with 250µL of APA766 stock) were grown in LB-kanamycin:DAP (50µg/mL kanamycin and 60µg/mL DAP). A 20mL culture was started from an individual *P. psychrophila* JB418 colony in LB broth and grown at RT overnight at 200 rpm. *E. coli* donor cells were washed twice with LB and resuspended in 25mL LB. Donor and recipient cells were then mixed at a 1:1 cell ratio based on OD600 measurements, pelleted, and resuspended in 100µL. This was done separately for each of the four *E. coli* cultures. 40µL were plated on nitrocellulose filters on LB plates with 60µg/mL DAP. Two filters were used for each of the four conjugation mixtures (8 total conjugations). The conjugations took place for 6 hours at RT. After 6 hours, the filters were each resuspended in 2mL of LB broth and then plated on LB:kanamycin (50µg/mL) for selection of transconjugants. 20 plates were plated of a 1:2 dilution for each conjugation (160 plates total). Transconjugants were pooled and harvested after three days of growth on selection plates. The pooled mixture was diluted back to 0.25 in 100mL of LB:kanamycin (50µg/mL). The culture was then grown at RT to an OD600 of 1.3 before glycerol was added to 10% final volume and 1mL aliquots were made and stored at −80°C for future use.

### TnSeq sequencing library preparation for *P. psychrophila* JB418 and TnSeq data analysis

Library preparation was performed as in Wetmore, et al. 2015 with slight modifications (Wetmore et al. 2015).

#### DNA extraction

DNA was extracted from the *P. psychrophila* JB418 RB-TnSeq library by phenol:chloroform extraction. Briefly, the cell pellet was vortexed at maximum speed for 3 minutes in the presence of 500µL buffer B (200mM NaCl,20mM EDTA sterilized by filtration), 210µL of 20% SDS, a 1:1 mixture of 425-600µM and 150-212µm acid-washed beads, and 500µL of phenol:chloroform, pH 8. The sample was then centrifuged for 3 minutes at 4°C at 8000 rpm prior to removing the aqueous phase to a new tube. 1/10 of sample aqueous phase volume of 3M sodium acetate was then added along with 1 aqueous phase volume of ice cold isopropanol. The sample was then placed for ten minutes at −80°C before centrifugation for five minutes at 4°C at 13000 rpm. The supernatant was discarded and 750µL of ice cold 70% ethanol was added before another centrifugation for five minutes at 4°C at 13000 rpm. The supernatant was discarded and the DNA pellet was allowed to air dry before resuspension in 50µL of nuclease-free water. DNA was quantified with Qubit double-stranded DNA high-sensitivity assay kit (Invitrogen, Carlsbad, CA).

#### DNA fragmentation and size selection

2µg of DNA was sheared with a Covaris E220 focused-ultrasonicator with the following settings: 10% duty cycle, intensity 5, 200 cycles per burst, 150 seconds. DNA was split into 2 aliquots (1µg each) and samples were size-selected for fragments of 300 bp using 0.85X Agencourt AMPure XP beads (Invitrogen) with a 1.4x ratio following the manufacturer’s instructions.

#### Library preparation

The entire 20µL volume of these two size-selected samples were then each used as input into the NEBNext End Prep step 1.1 of the NEBNext Ultra DNA Library Prep Kit for Illumina (New England Biolabs, Ipswich, MA) protocol. The remainder of the manufacturer’s protocol was then followed with the exception that for adapter ligation, we used 0.8µL of 15µM double-stranded Y adapters. Adapters were prepared by first combining 5µL of 100µM Mod2_TS_Univ and 5µL of 100µM Mod2_TruSeq. This mixture was then incubated in a thermocycler for 30 min at 37°C, followed by ramping at 0.5°C per second to 97.5°C before a hold at 97.5°C for 155 seconds. The temperature was then decreased by 0.1°C per five seconds for 775 cycles, followed by a hold at 4°C. Annealed adapters were diluted to 15µM in TE and stored at −80°C before use. AMPure XP ratios for a 200bp insert size were used as recommended in Table 1.1 of the NEBNext Ultra DNA Library Prep Kit for Illumina manual.

To enrich for transposon-insertion sites, PCR amplification was done on the adapter-ligated DNA with NEBNext Q5 Hot Start HiFi Master Mix and Nspacer_barseq_pHIMAR and P7_MOD_TS_index3 primers (Wetmore et al. 2015) with the following program: 98°C 30 sec, 98°C 10 sec, 65°C 75 sec, repeat steps 2-3 24X, 65°C 5 min, and then maintained at 4°C. Following PCR and clean-up of step 1.5 of the NEBNext Ultra DNA Library Prep Kit for Illumina manual, the two preps were pooled and the concentration was quantified with Qubit double-stranded DNA high-sensitivity assay kit (Invitrogen). A second size selection clean-up was performed by repeating step 1.5 of the NEBNext Ultra DNA Library Prep Kit for Illumina manual.

#### Library sequencing

The sample was analyzed on an Agilent TapeStation and the average size was 380bp and the concentration was 57pg/µL. This sample was then submitted for sequencing on a HiSeq 2500 Rapid Run (150bp fragments, paired-end) at the UCSD IGM Genomics Center:

#### Library characterization

TnSeq reads were analyzed with the perl script MapTnSeq.pl from (Wetmore et al. 2015). That script maps each read to the *P. psychrophila* genome. The script DesignRandomPool.pl (Wetmore et al. 2015) was used to generate the file containing the list of barcodes that consistently map to a unique location as well as their location.

### RB-TnSeq experiments for *E. coli* and *P. psychrophila* JB418

The *E. coli* barcoded transposon library Keio_ML9 and the *P. psychrophila* strain JB418 library were used for fitness assays on CCA during growth alone, growth in pairwise condition with each bloomy rind cheese community member and during growth with the full community.

#### Library pre-culture

Each library has to be initially amplified before use. One aliquot of each library was thawed and inoculated into 25mL of liquid LB-kanamycin (50µg/mL). Once the culture reached mid-log phase (OD=0.6-0.8), 5mL of that pre-culture was pelleted and stored as the T0 reference for the fitness analysis. The remaining cells were used to inoculate the different fitness assay conditions.

#### Inoculations

For each fitness assay, 7*10^6^ cells of the library pre-culture were inoculated on 10% cheese curd agar, pH 7 after having been washed in PBS1x-Tween0.05%. For each pairwise assay, 7*10^6^ cells of the partner were co-inoculated with the library. For the community assay, 7*10^6^ cells of H. lavei JB232 and *G. candidum* as well as 7*10^5^ cells of *P. camemberti* were co-inoculated with the library. For each condition, assays were performed in triplicates.

#### Harvest for gene fitness calculation

Harvests were performed at T= 24h, 48h and 72h. Sampling was done by flooding a plate with 1.5mL of PBS1X-Tween0.05% and gently scraping the cells off. The liquid was then transferred into a 1.5mL microfuge tube and cells were pelleted by centrifugation. After removing the supernatant, the cells were washed in 1 mL of RNA-protect solution (Qiagen, Hilden, Germany), pelleted and stored at −80°C before further experiments.

#### gDNA and mRNA extraction

gDNA and mRNA were simultaneously extracted by a phenol-chloroform extraction (pH 8) from samples of the competitive assays. For each extraction: 125µL of 425-600µm acid-washed beads and 125µL of 150-212µm acid-washed beads were poured in a screw-caped 2mL tube. 500µL of 2X buffer B (200mM NaCl, 20mM EDTA) and 210µL of SDS 20% were added to the tube as well as the pellet and 500µL of Phenol:Chloroform (pH 8). Cells were lysed by vortexing the tubes for 2 minutes at maximum speed. Aqueous and organic phases were separated by centrifugation at 4°C, 8KRPM for 3 minutes and 450µL of the aqueous phase (upper phase) was recovered in a 1.5mL eppendorf tube. 45µL of Sodium Acetate 3M and 450µL of ice cold isopropanol were added before incubating the tubes at −80°C for 10 minutes. The tubes were then centrifuged for 5 minutes at 4°C at 13KRPM. The pellet was then washed in 750µL of 70% ice cold ethanol and re-suspended in 50µL of DNAse/RNAse free water. Each sample was split into 2 times 25µL and stored at −80°C until further analysis.

#### Library preparation and sequencing

After gDNA extraction, the 98°C BarSeq PCR as described and suggested in Wetmore et al., 2015 was used to amplify only the barcoded region of the transposons. Briefly, PCR was performed in a final volume of 50µL: 25µL of Q5 polymerase master mix (New England Biolab), 10µL of GC enhancer buffer (New England Biolab), 2.5µL of the common reverse primer (BarSeq_P1 – Wetmore et al., 2015) at 10µM, 2.5µL of a forward primer from the 96 forwards primers (BarSeq_P2_ITXXX) at 10µM and 50ng to 2µg of gDNA. For each triplicate, the PCR was performed with the same forward primer so all replicates of a condition could be pooled and have the same sequencing multiplexing index. So, for *E. coli* analysis, we performed 46 PCRs (1 T0 samples and 45 harvest samples) involving 16 different multiplexing indexes. For *P. psychrophila* JB418 analysis, we performed 46 PCR (1 T0 and 45 harvest samples) involving 16 other multiplexing indexes. We used the following PCR program: (i) 98°C - 4 minutes, (ii) 30 cycles of: 98°C – 30 seconds; 55°C – 30 seconds; 72°C – 30 seconds, (iii) 72°C – 5 minutes. After the PCR, 10µL of each of the 92 PCR products were pooled together to create the BarSeq library(920µL) and 200µL of the pooled library were purified using the MinElute purification kit (Qiagen). The final elution of the BarSeq library was performed in 30µL in DNAse and RNAse free water.

The BarSeq library was then quantified using Qubit® dsDNA HS assay kit (Invitrogen) and sequenced on HiSeq4000 (50bp, single-end reads), by the IGM Genomics Center at the University of California, San Diego.

#### Data processing and fitness analysis

BarSeq data processing and gene fitness calculation was performed separately for the *E. coli* and the *P. psychrophila* JB418 experiments. For each library, BarSeq reads were processed using the custom perl script BarSeqTest.pl (Wetmore et al. 2015). This script combines two perl scripts essential for the BarSeq data processing. After the raw reads have been de-multiplexed into each conditions, the computational pipeline: (i) identifies each barcode and the associated number of reads, (ii) calculates the strain fitness for each insertion mutant as the log2 ratio of counts in the condition and in the T0 sample and (iii) calculates each “gene fitness” as the average of strain fitness of all the insertion mutants associated with that gene as well as calculation of a fitness t-score. The following rules were applied during the fitness calculations: (i) only insertion mutants with barcode insertion within the 10%-90% fraction of the genes were considered, (ii) barcodes with less than 3 reads in the T0 were ignored and (iii) genes with less than 30 counts in T0 were ignored. For each library, the pipeline uses a table where each barcode is mapped to a location in the genome. The Arkin lab (Physical Biosciences Division, Lawrence Berkeley National Laboratory, Berkeley, California, USA) kindly provided the TNSeq table for the *E. coli* library and we generated a TNSeq table for *P. psychrophila* strain JB418.

We calculated *E. coli* and *P. psychrophila* JB418 genes fitnesses at T=24h, 48h and 72h in the following conditions: growth alone, growth with *H. alvei*, growth with *G. candidum*, growth with *P. camemberti* and growth with the community.

### Keio collection mutant competitive assays

We used mutants from the Keio collection to validate the genes identified by RB-TNseq as having a significant fitness in *E. coli* (see list in supplementary data S3). Each mutant was grown in a competition assay with the wild type (Keio ME9062 –(Baba et al. 2006)). 1000 cells of a specific mutant were inoculated with 1000 cells of the wild type on the surface of the same cheese plug in a 96 well plate containing 10% cheese curd agar, pH7. The number of the mutant cells and the WT cells were calculated at T0 and day 1 after harvesting, homogenizing the cheese plug, plating serial dilutions and counting CFUs. Experimental fitness of each mutant was calculated as the log2 of the mutant abundance (mutant CFUs divided by total CFUs (WT + mutant)) after 24 hours and its abundance at T0.

### RNASeq and differential expression analysis

The RNASeq analysis was performed for the *E. coli* experiments only.

#### RNASeq libraries preparations

Libraries were prepared for duplicates of the following conditions: *E. coli* growth alone, with *H. alvei*, with *G. candidum*, with *P. camemberti* and with the community for T= 24h, 48h and 72h. RNA samples from the *E. coli* BarSeq experiment were used to produce the RNASeq library.

Each library was prepared as follows. First, RNA samples were treated with DNAse using the “Rigorous DNAse treatment” for the Turbi DNA-free kit (AMBIO – Life Technologies, USA) and RNA concentration was measured by nucleic acid quantification in Epoch Microplate Spectrophotometer (BioTek, Winooski, VT). Then, transfer RNAs and 5S RNA were removed using the MEGAclear Kit Purification for Large Scale Transcription Reactions (AMBION, Life Technologies, Waltham, MA) following manufacturer instructions. Absence of tRNA and 5S RNA was verified by running 100ng of RNA on a 1.5% agarose gel and RNA concentration was quantified by nucleic acid quantification in Epoch Microplate Spectrophotometer. Also, presence of trace amounts of genomic DNA was assessed by PCR using universal bacterial 16S PCR primers (Forward primer: AGAGTTTGATCCTGGCTCAG, Reverse Primer: GGTTACCTTGTTACGACTT). Final volume of the PCR was 20µL: 10µL of Q5 polymerase master mix (New England Biolab), 0.5µL of forward primer 10uM, 0.5µL of reverse primer 10uM and 5µL of non-diluted RNA. PCR products were run on a 1.7% agarose gel and when genomic DNA was amplified, another DNAse treatment was performed as well as a new verification of absence of genomic DNA. Ribomosomal RNA depletion was performed using the Ribo-Zero rRNA removal kit by Illumina (Illumina, San Diego, CA). Ribosomoal RNA depletion was performed according to manufacturer instructions, we used 1µL of RiboGuard RNAse Inhibitor in each sample as suggested, followed instructions for 1-2.5ug of RNA input and we used a 2:1 mix of bacterial Ribo-Zero Removal solution and yeast Ribo-Zero Removal solution. rRNA depleted samples were recovered in 10µL after ethanol precipitation. Concentrations after ribodepletion were measured using Qubit® RNA HS Assay Kits (Invitrogen). The RNASeq library was produced using the NEBNext®UltraTM RNA Library Prep Kit for Illumina® for purified mRNA or ribosome depleted RNA. We prepared a library with a fragment size of 300 nucleotides and used the 10uM NEBNext Multiplex Oligos for Illumina (Set 1, NEB #E7335) lot 0091412 and the NEBNext multiplex Oligos for Illumina (Set 2, NEB #E7500) lot 0071412. We did the PCR product purification with 0.8X Agencourt AMPure XP Beads instead of 0.9X. Library samples were quantified with Qubit® DNA HS Assay Kits before the quality and fragment size were validated by TapeStation (HiSensD1000 ScreenTape). Library samples were pooled at a concentration of 15nM each and were sequenced on HISeq4000 (50bp, single-end).

#### Differential expression analysis

RNASeq reads were mapped to the concatenated genome of *Escherichia coli* K12 BW25113 (Grenier et al. 2014) and *H. alvei* using Geneious version R9 9.1.3 (http://www.geneious.com, (Kearse et al. 2012))). Only the reads that uniquely mapped to a single location on the *E. coli* genome section were conserved. *E. coli* and *H. alvei* genome are divergent enough so 50 nucleotides reads potentially originating from *H. alvei* mRNA would not map to *E. coli* genome and few reads from *E. coli* would map on *H. alvei* genome.

*E. coli* expression analysis was performed using the R packages: Rsamtool (R package version 1.30.0), GenomeInfoDb (R package version 1.14.0.), GenomicFeatures (Lawrence et al. 2013), GenomicAlignments, GenomicRanges (Lawrence et al. 2013) and DESeq2 (Love et al. 2015). We followed the workflow described by Love et al. and performed the differential expression analysis using the package DESeq2. Differentially expressed genes between two conditions were selected with an adjusted p-value lower than 1% and an absolute log of fold change equal or greater than 1.

### KEGG pathway enrichment analysis

Functional enrichment analysis was performed using the R package clusterProfiler (Yu et al. 2012). We used the latest version of the package org.EcK12.eg.db for *E. coli* annotations (R package version 3.5.0.). We used Benjamini-Hochberg for multiple comparison correction and only the KEGG pathways enriched with an adjusted p-value lower than 5% were considered. The *E. coli* database was used with the simplify function to avoid redundant GO Term and only display the GO Term with the highest significance within a cluster.

## ACKNOWLEDGMENTS

The authors would like to thank: the Arkin lab and the Deutschbauer lab at UC Berkeley/LBNL for the *E. coli* Keio_ML9 library, Morgan Price for guidance on RB-TNseq analysis, Steven Villareal and Tyler Nelson for technical assistance, Kristen Jepsen at the IGM Genomics Center at UC San Diego for assistance with sequencing, Ben Wolfe and Sandeep Venkataram for their constructive comments on this work as well as all the Dutton lab members for their input.

---

**Supplementary figure 1:**
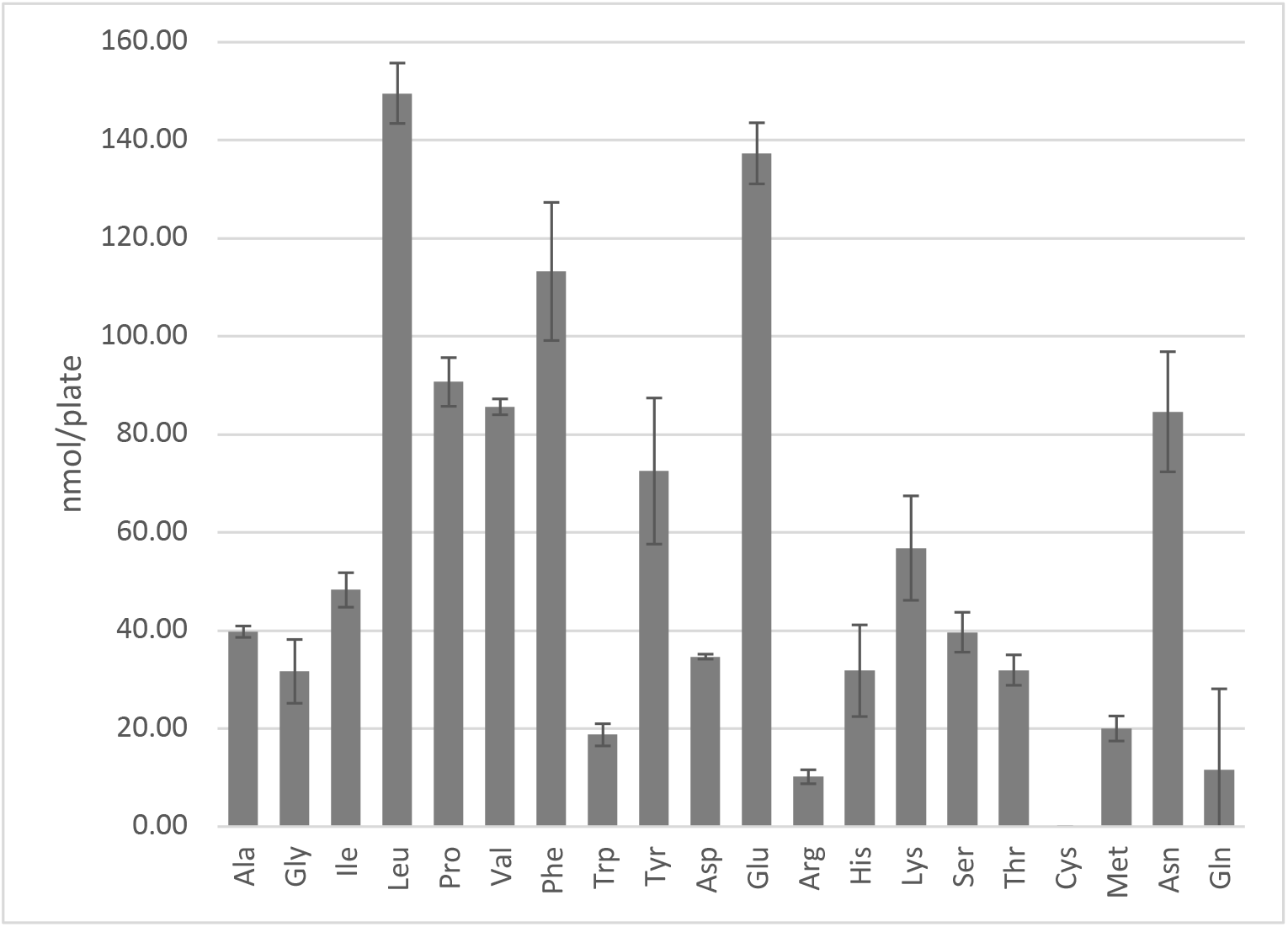
Quantification of free amino-acids in CCA. Free amino acid characterization and quantification have been carried out by the Proteomics & Mass Spectrometry Facility of the Donald Danforth Plant Science Center and each analysis has been performed in triplicate. Samples were prepared as according to the Proteomics & Mass Spectrometry Facility of the Donald Danforth Plant Science Center instructions. For free amino acid analysis 150 mg CCA were frozen in liquid nitrogen and grinded in the presence of 600 μL of water:chloroform:methanol (3:5:12 v/v). Tubes were then centrifugated at full speed for two minutes and supernatant was recovered in a new 2mL eppendorf tube. A second extraction with 600uL of water:chloroform:methanol was performed followed by a two minute centrifugation at full speed. Supernatant was then recovered and combined with the previous supernatant in a 2mL eppendorf tube. Then 300uL of chloroform and 450uL of water were added before centrifugation at full speed for two minutes. The upper phase was recovered and transferred to a new tube. Samples were dried in a speedvac overnight and stored at −20°C. The total concentration of free amino acids in CCA is 75.3 nmol/mL. Analysis of total amino acids was also performed by the Proteomics & Mass Spectrometry Facility of the Donald Danforth Plant Science Center. It highlights that free aminos are a very small fraction of total amino acid whose concentration is 16.5 ±2,97 μmol/mL, six times less than the LB concentration measured by Sezonov et al., 2012. Ala: alanine, Gly: glycine, Ile: isoleucine, Leu: leucine, Pro: proline, Val: valine, Phe: phenylalanine, Trp: tryptophame, Tyr: tyrosine, Asp: aspartate, Glu: glutamate, Arg: arginine, His: histidine, Lys: lysine, Ser: serine, Thr: threonine, Cys: cysteine, Met: methionine, Asn: asparagine, Gln: glutamine.

**Supplementary figure 2:**
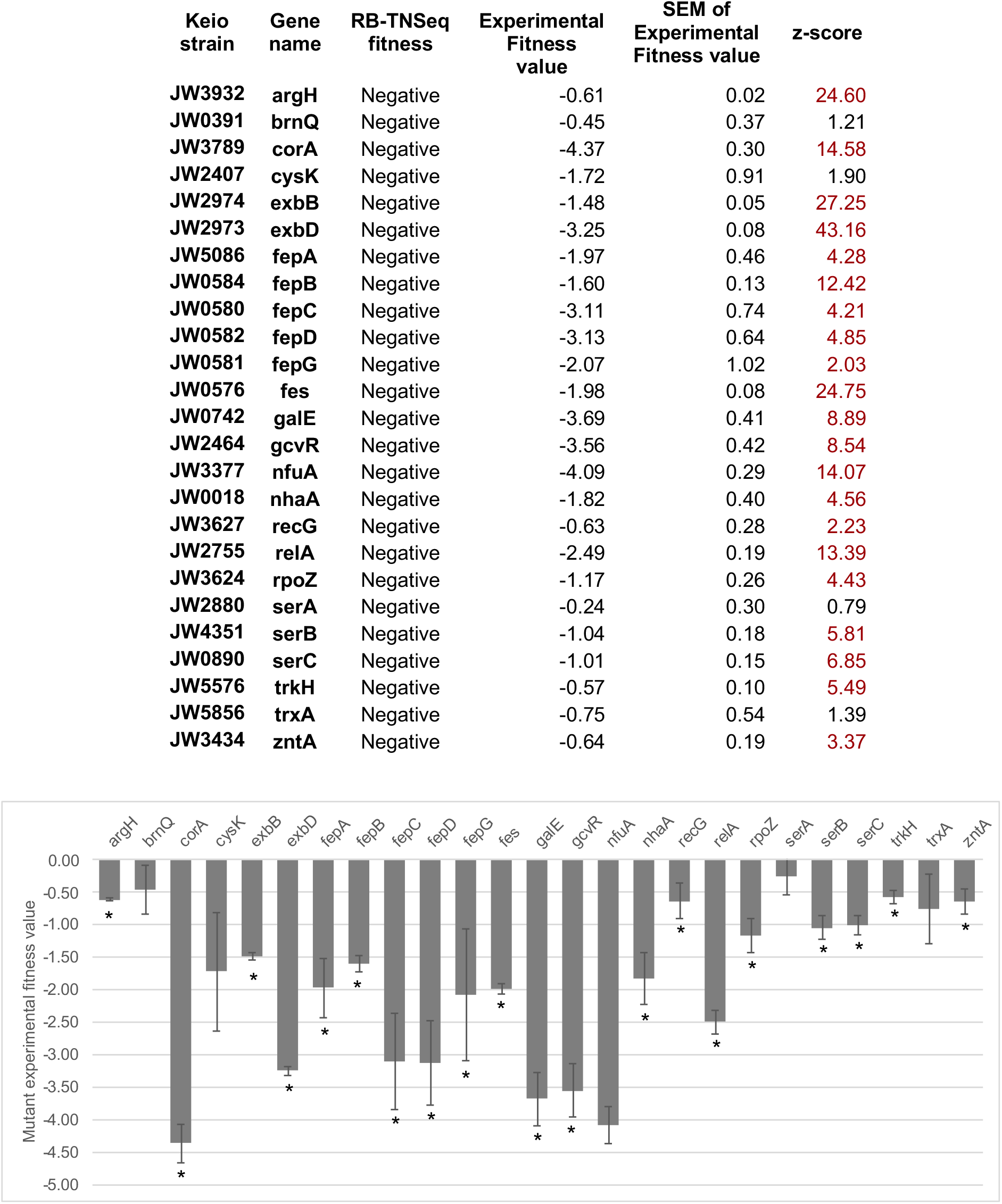
Competitive assays of 35 mutants of the Keio collection (Baba et al., 2006). Competition assays between single knock outs and the wild-type strain have been carried out for 35 strains associated with genes identified as essential for *E. coli* growth using RB-TnSeq (Significant fitness lower than −1 after 1 day of growth). * highlights fitness values different from 0 with a confidence higher than 95%

**Supplementary figure 3:**
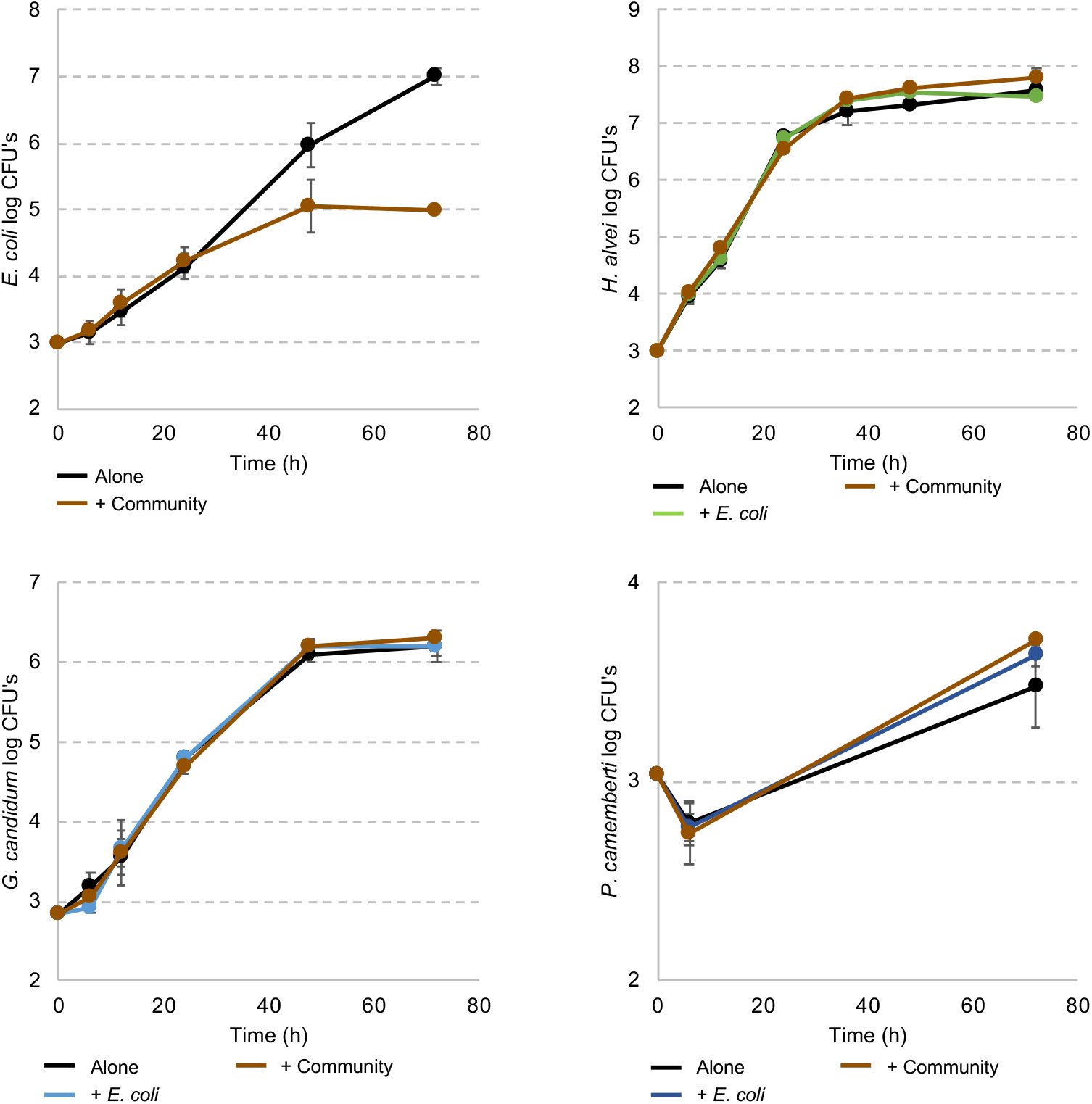
*E. coli* and community member growth curves alone, in pairwise conditions or during community growth. Each graph represents the growth over time of *E. coli*, *H. alvei*, *G. candidum* or *P. camameberti* alone, in pairwise growth or with the community.

**Supplementary figure 4:**
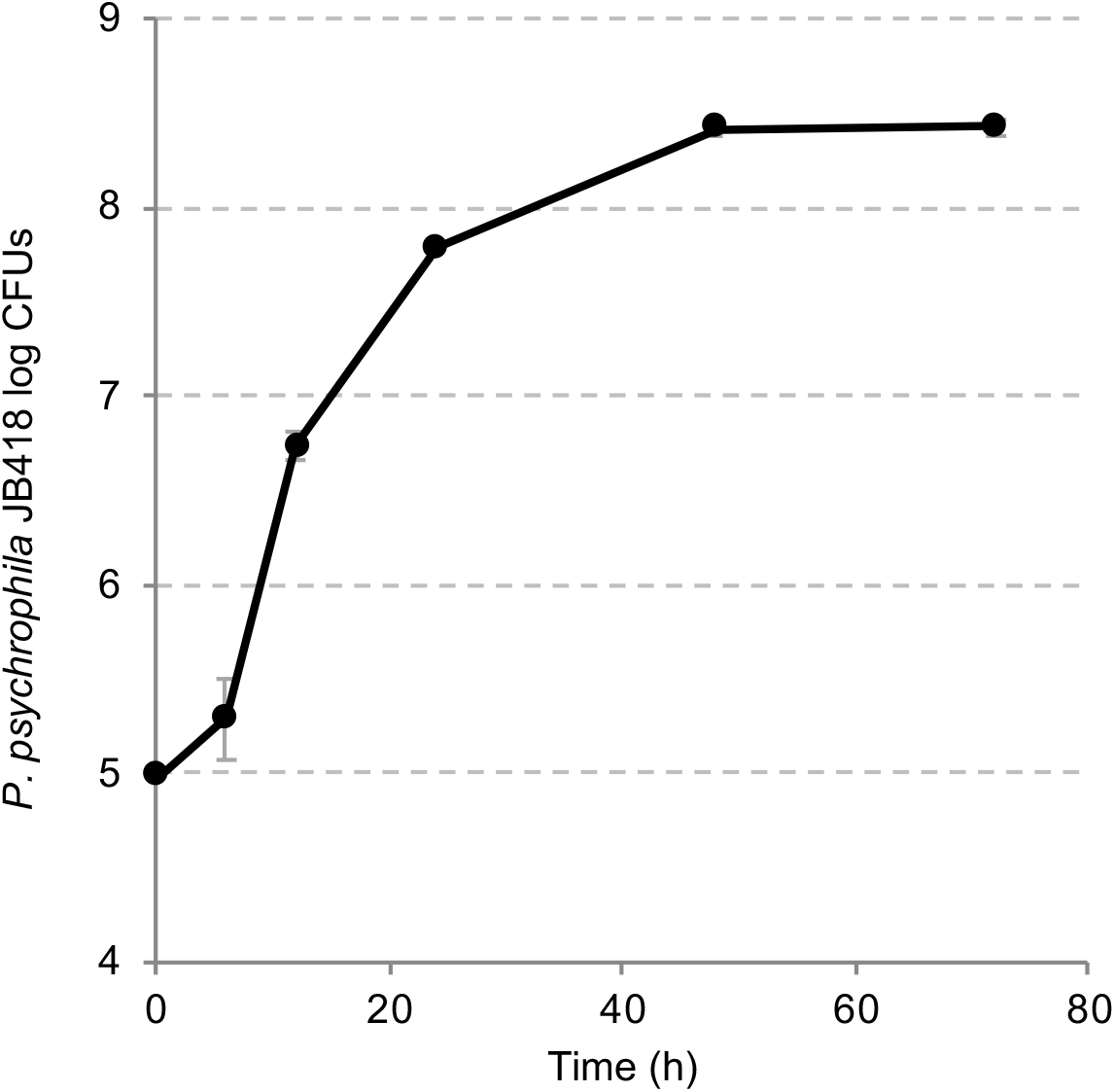
*P. psychrophila* JB418 growth curve on CCA.

**Supplementary figure 5:**
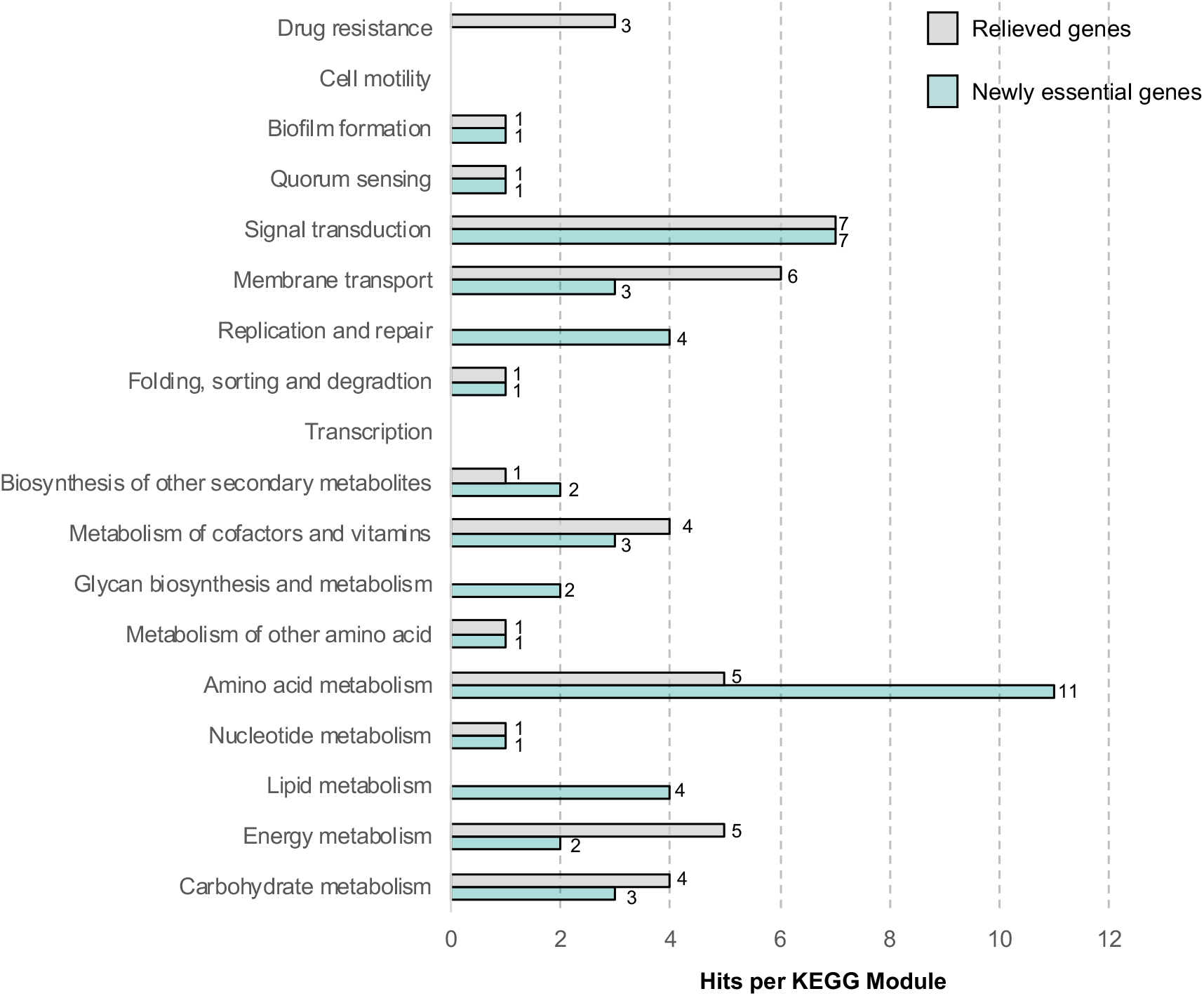
Functional analysis of newly essential genes and relieved genes for *P. psychrophila* JB418 growth in pairwise condition versus alone.

